# An evolution-based framework for describing human gut bacteria

**DOI:** 10.1101/2023.12.04.569969

**Authors:** Benjamin A. Doran, Robert Y. Chen, Hannah Giba, Vivek Behera, Bidisha Barat, Anitha Sundararajan, Huaiying Lin, Ashley Sidebottom, Eric G. Pamer, Arjun S. Raman

## Abstract

The human gut microbiome contains many bacterial strains of the same species (‘strain-level variants’). Describing strains in a biologically meaningful way rather than purely taxonomically is an important goal but challenging due to the genetic complexity of strain-level variation. Here, we measured patterns of co-evolution across >7,000 strains spanning the bacterial tree-of-life. Using these patterns as a prior for studying hundreds of gut commensal strains that we isolated, sequenced, and metabolically profiled revealed widespread structure beneath the phylogenetic level of species. Defining strains by their co-evolutionary signatures enabled predicting their metabolic phenotypes and engineering consortia from strain genome content alone. Our findings demonstrate a biologically relevant organization to strain-level variation and motivate a new schema for describing bacterial strains based on their evolutionary history.

**One Sentence Summary:** Describing bacterial strains in the human gut by a statistical model that captures their evolutionary history provides insight into their biology.

## Main

Microbial communities (‘microbiomes’) are ubiquitous across diverse environments, spanning oceans to individual humans (*1–4*). One microbiome relevant to human health is the gut microbiome: the trillions of microorganisms residing along the intestinal tract of humans (*5*, *6*). A number of studies have demonstrated the significance of the gut microbiota—the bacteria within the microbiome—for influencing host physiology and predilection for developing several diseases (*7*). This has led to many efforts describing the composition of human gut microbiotas and understanding how composition affects microbiome function. As interrogating microbiomes has become easier, the incredible taxonomic complexity of microbiotas and associated functional consequences have become more appreciated than ever before (*8*, *9*).

Studies focused on cataloguing microbiota composition have revealed the extensive presence of ‘strain-level variants’: strains that belong to the same species but are genetically different (*10–12*). Moreover, several case studies have highlighted the direct impact of individual strains on gut microbiome function and host health. For instance, reconstitution of the infant gut microbiota using *Bifidobacterium longum* subspecies *infantis*—a strain of *B. longum* that metabolizes human milk oligosaccharides—has been shown to repair intestinal inflammation due to acute malnutrition in humans (*13–15*). In another example, profiling different strains of *Bacteroides ovatus* showed differential capacity in inducing immunoglobulin A levels (*16*). With respect to influence on bacterial fitness in the gut, interrogation of *Bacteroides* and *Parabacteroides* strains revealed strain-level preferences for binding polysaccharides (*17*). Many similar vignettes, from understanding the etiology of food-borne outbreaks to characterizing the repair of the gut microbiota following antibiotic exposure, have highlighted the functional role of strain-level variants (*18*, *19*).

As the importance of individual bacterial strains has become increasingly appreciated, an outstanding question is how should strains be described? If considering complete genomes at amino acid resolution, almost every newly procured strain is a ‘strain-level variant’ because it is likely to be different in some way from other strains of the same species. If instead the whole genome is compressed into a more practically manageable description, like the 16S rDNA sequence or sets of marker genes, strains become collapsed into their phylogenetic description obscuring potentially important adaptive changes. With respect to classifying strains by biological function, recent studies that created banks of sequenced and phenotyped gut bacterial strains have shown a common trend: phenotypic differences between strains of the same species are difficult to understand (*20–22*). As an example, it has been shown that metabolic capacities of bacteria follow coarse phylogeny but that variation amongst individual strains is mostly unrelated to metabolic variability (*20*). Collectively, these observations have led to defining strain-level variants taxonomically, detailing differences between strains at increasing molecular resolution, rather than conceptually where strains are distinguished by biologically relevant differences (*23*, *24*). Thus, the status quo strategy is to functionally interrogate each and every new strain because structure amongst strain-level variants, i.e. ‘subspecies phylogeny’, is difficult to ascertain and unlikely to be associated with strain-level phenotype.

A key limitation of performing comparative analysis on strain-level variants within strain banks is the tremendous degree of phylogenetic under-sampling compared to the bacterial tree-of-life. Strain banks usually contain strains from a specific econiche—only from the human gut for instance—and therefore reflect a small portion of phylogenetic diversity. This limitation skews our understanding of gene content that is under selective pressure versus gene content that is allowed to vary. Establishing a relationship between strain genome and function through comparative analysis therefore becomes entangled with genomic variation associated with phylogeny (*25*).

The existence of large databases of sequenced bacteria motivated a hypothesis that we tested here. Namely, that by leveraging the vast diversity of sequenced strains procured from many different environments, we could ‘fill in’ evolutionary distance within a given strain bank in a more complete way. That is, constraints gleaned from the evolutionary record could be used as a Bayesian prior for contextualizing differences between strains procured form a single environment—the human gut (*26*, *27*). If achieved, this space of evolutionary relationships may resolve fine-grained differences between strains of the same species and would allow testing whether phenotypic qualities specific to strain-level variants could be learned from genetic information.

We created a strain bank of 669 gut commensal strains that were isolated from fecal samples collected across 28 healthy human donors, whole-genome sequenced, and metabolically profiled. Consistent with previous studies, traditional analysis of this strain bank from 16S rDNA sequence and well-known sets of marker genes could not resolve genomic differences between strain-level variants or their associated metabolic qualities. We next developed a new statistical framework for inferring evolutionary distance between bacteria based on patterns of co-evolution. We used this framework to create a tree of bacterial relationships across >7,000 non-redundant bacterial proteomes collected across many diverse environments that we termed a ‘Spectral Tree’. Examining our strain bank within the structure of the Spectral Tree revealed widespread subspecies phylogeny across gut commensal strains. Interrogation of the Spectral Tree revealed that subspecies phylogeny was driven by a history of host phage exposure amongst groups of donors and was associated with a loss of well-conserved bacterial genetic machinery. We created a new descriptor for a given bacterial strain based on its unique hierarchical sequence of branching paths in the Spectral Tree. Training a simple LASSO regression on the pattern of Spectral Tree branches assigned to each strain accurately predicted strain-level metabolic output deviating from general phylogenetic trends for key gut-related metabolites. Applying our LASSO models to newly acquired strains outside our strain bank enabled rationally engineering consortia with desired metabolic traits using only the genomes of individual strains. Together, our results show that describing bacterial strains by their co-evolutionary signatures is a natural, data-driven, interpretable, and biologically useful representation of strain genomes.

## Results

### A bank of 669 human gut commensal strains

We isolated and sequenced over 1,000 commensal bacterial strains from the feces of 28 healthy human volunteers. Our resulting bank of gut commensal strains (‘commensal strain bank’ from here on) comprised 669 diverse strains that we whole genome sequenced (**Fig. 1A, Table S1**) (Methods). The commensal strain bank was enriched for gram negative anaerobes within *Lachnospiraceae*, *Bacteroidaceae*, and *Bifidobacteriaceae* families (**Fig. 1B**). We metabolically profiled all strains within the strain bank across 50 targeted metabolic features comprising amino acids, aromatics, branch-chained fatty acids, indoles, phenolic aromatics, and short-chained fatty acids (Methods). Of the 50 features, 32 showed detectable changes in relative abundance across at least 10% of strains. For these 32 features, we found there was considerable strain-level metabolic variability across numerous species in the strain bank (**Fig. S1**, **Table S2**).

**Fig. 1.**
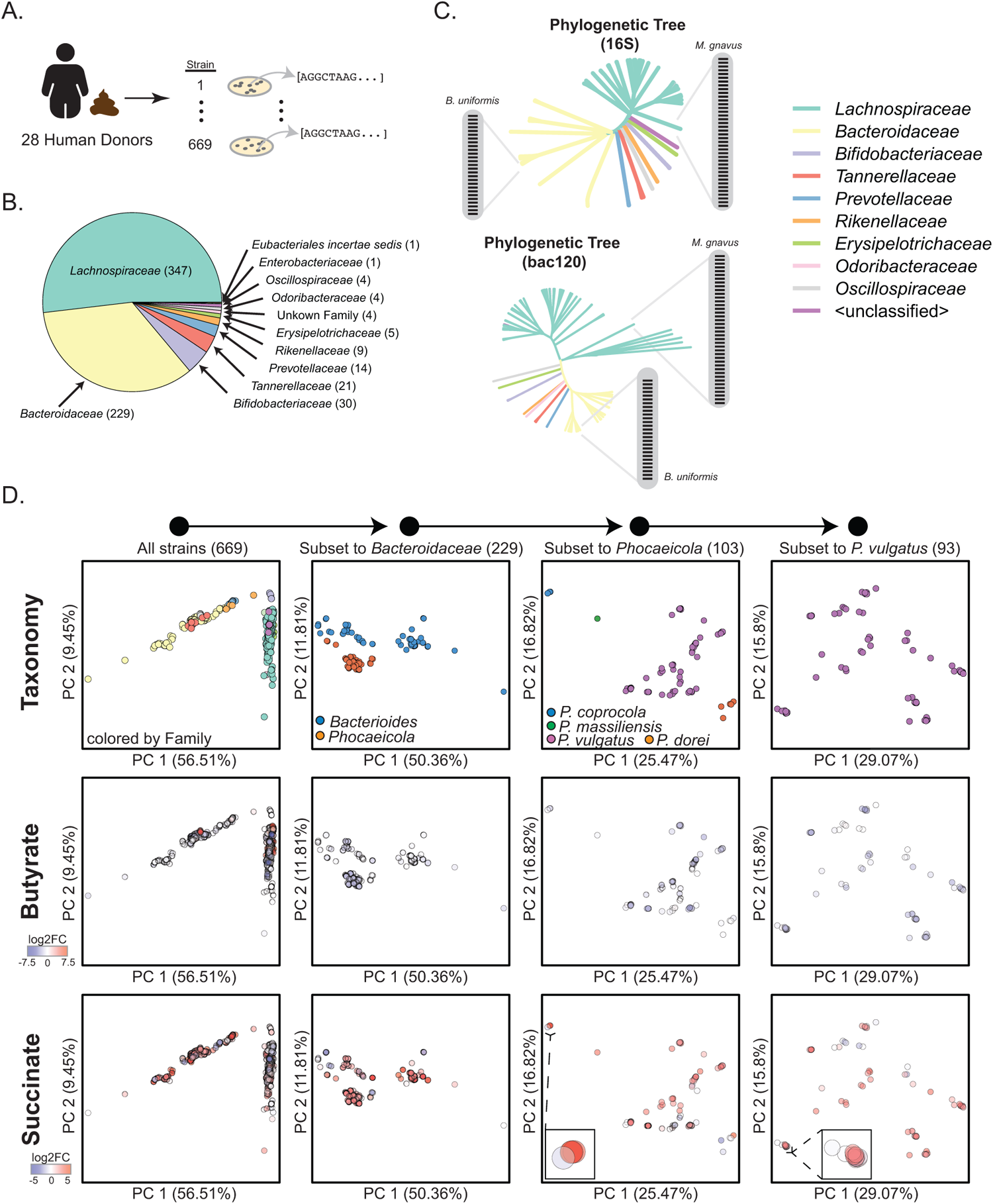
Establishing and characterizing a bank of commensal bacterial strains from the human gut. **(A)** 669 gut commensal strains were isolated, cultured, and whole-genome sequenced from fecal samples of 28 healthy human donors. **(B)** Phylogenetic distribution of commensal strain bank; number of strains belonging to specific phylogenetic family in parenthesis. **(C)** Phylogenetic trees of commensal strain bank defined by either 16S rDNA sequence (upper panel) or set of 120 genes conserved across the bacterial tree-of-life (‘bac120’, lower panel). Insets show that both phylogenetic trees do not resolve strain-level variation amongst *Mediterraneibacter gnavus* or *Bacteroides uniformis* strains. **(D)** PCA plots of strains defined by orthologous gene groups (OGGs). Each dot in the PCA plots is a strain. From left to right, commensal strain bank is subset to strain-level variation amongst *Phocaceicola vulgatus* strains. The top row is colored by phylogeny, the middle and bottom rows are colored by relative changes in the concentrations of butyrate (middle) and succinate (bottom) for each strain.

We created phylogenetic trees of our strain bank defined by (i) the 16S rDNA sequence and (ii) 120 proteins used to create phylogenetic relationships defined by the GTDB database (‘bac120’) (Methods) (*28*). We found that both phylogenetic trees robustly defined coarse phylogenetic differences but were limited in their capacity to resolve differences between strains belonging to the same species (**Fig. 1C**). To consider a wider range of the bacterial genome than the 16S region or the bac120 proteins, we annotated all strains by their orthologous gene group (OGG) content and conducted principal components analysis (PCA) as well as a Uniform Manifold Approximation (UMAP) based analysis of the resulting alignment (Methods) (*29*, *30*). Like the phylogenetic trees, PCA and UMAP strain clusters were resolved by coarse phylogenetic differences but did not resolve strain-level variation (**Fig. S2**). Moreover, the relative placement of strains did not correlate with their metabolic similarity (**Fig. 1D**) (Methods). Together, these results were consistent with previously published studies illustrating the difficulty in distinguishing strains of the same species by both genome and function.

### A measure of evolutionary distance based on patterns of bacterial co-evolution

To test our hypothesis that evolutionary relationships across a wealth of sequenced bacteria could aid in revealing strain-level genomic differences within our strain bank, we turned to a large database of sequenced non-redundant bacterial strains procured across a diversity of environments. Our previous work described a phylogenomic analysis of the kingdom Bacteria using all reference proteomes in the UniProt database (>7,000 strains in total) (*31*, *32*). Analysis of this database illustrated that co-evolutionary patterns defined a hierarchy of phylogeny. Major principal components clustered bacteria belonging to the same phylum, deeper components class, and so on until species. Building upon this finding, we reasoned that the whole principal component spectrum of bacterial co-evolution defined across the UniProt database, including principal components typically discarded as noise, may be useful for inferring evolutionary distances between bacteria and resolve fine-grained differences between strains of the same species.

Developing a definition of evolutionary distance inferred from patterns of bacterial co-evolution first required studying simple, manageable models of the evolutionary process. Thus, we used ‘toy model’ simulations of diversification and selection to explore how principal components could be used to define an evolutionary distance metric between taxa that are defined by a complex set of features, like a genome. Our mathematical and computational workflow is described in detail within the Supplementary Text (Supplementary Text, Sections 1 and 2). Briefly, given an alignment of taxa, we found that the generations of sequential diversifications giving rise to taxa were directly related to patterns of co-variation between taxa. This finding enabled inferring a tree of taxonomic relatedness that accurately captured the precedent sequence of diversification events (**Fig. S3-S5**).

While for simple trajectories of sequential diversification our approach matched results of existing tree-based inference methods, our approach was uniquely suited for addressing our hypothesis in three ways. First, for complex trajectories such as convergent processes, our approach resolved paths of diversification while existing maximum-likelihood and Bayesian approaches resulted in erroneous placement of taxa (**Fig. S6**) (Supplementary Text, Section 3). Second our approach required substantially less computational resources than existing methods for building a tree of taxonomic relationships (**Fig. S7**) (Methods). This quality made inferring a tree across thousands of bacteria, like the UniProt database, a tractable goal that is not feasible with current approaches. Finally, because our approach was based on straightforward matrix calculations, we could infer the evolutionary distance of newly acquired strains within the context of a large set of previously characterized taxa. Thus, collections of new strains like our strain bank could be ‘projected into’ our tree thereby making our tree a dynamic object capable of incorporating more sequences to reflect the increasing corpus of data.

As our approach infers a tree of relatedness from principal components of taxonomic covariation, we term the tree resulting from our approach a ‘Spectral Tree’. We constructed a Spectral Tree for the set of non-redundant bacterial proteomes in UniProt (Supplementary Text, Section 4) (Methods). We found that the topology of the tree matched a hierarchy of phylogeny: shallow to deep Spectral Tree clusters progressively grouped bacteria from the same phylum, class, order, family, genus, and species (**Fig. S8**). Additionally, interrogation of specific bacterial Families common to the gut microbiota illustrated clustering consistent with canonical taxonomic classifications (**Fig. S9**). These results illustrated that the Spectral Tree captured known evolutionary relationships amongst bacteria.

### Discovering subspecies phylogeny in our strain bank

We used the Spectral Tree constructed from UniProt strains to study our commensal strain bank. We annotated all strains in our strain bank by their OGG content and projected each strain into the Spectral Tree (**Fig. 2A, Figs. S10-S11**) (Methods). We compared the distances of all pairs of strains in our strain bank that share the same genus or species designations computed from (i) the Spectral Tree, (ii) the phylogenetic tree created from the 16S rDNA sequence, or (iii) the phylogenetic tree created from the amino acid sequence of bac120. We found that for pairs of strains from the same genus or species, the Spectral Tree resolved differences between our strains more than the other phylogenetic trees to a statistically significant degree (**Fig. 2B**). The average relative distance of strain pairs based on 16S- and bac120 trees was nearly zero; while the same distribution based on the Spectral Tree was bimodal (**Fig. 2B**, inset). This result suggested that the Spectral Tree was resolving subspecies-level differences between strains.

**Fig. 2.**
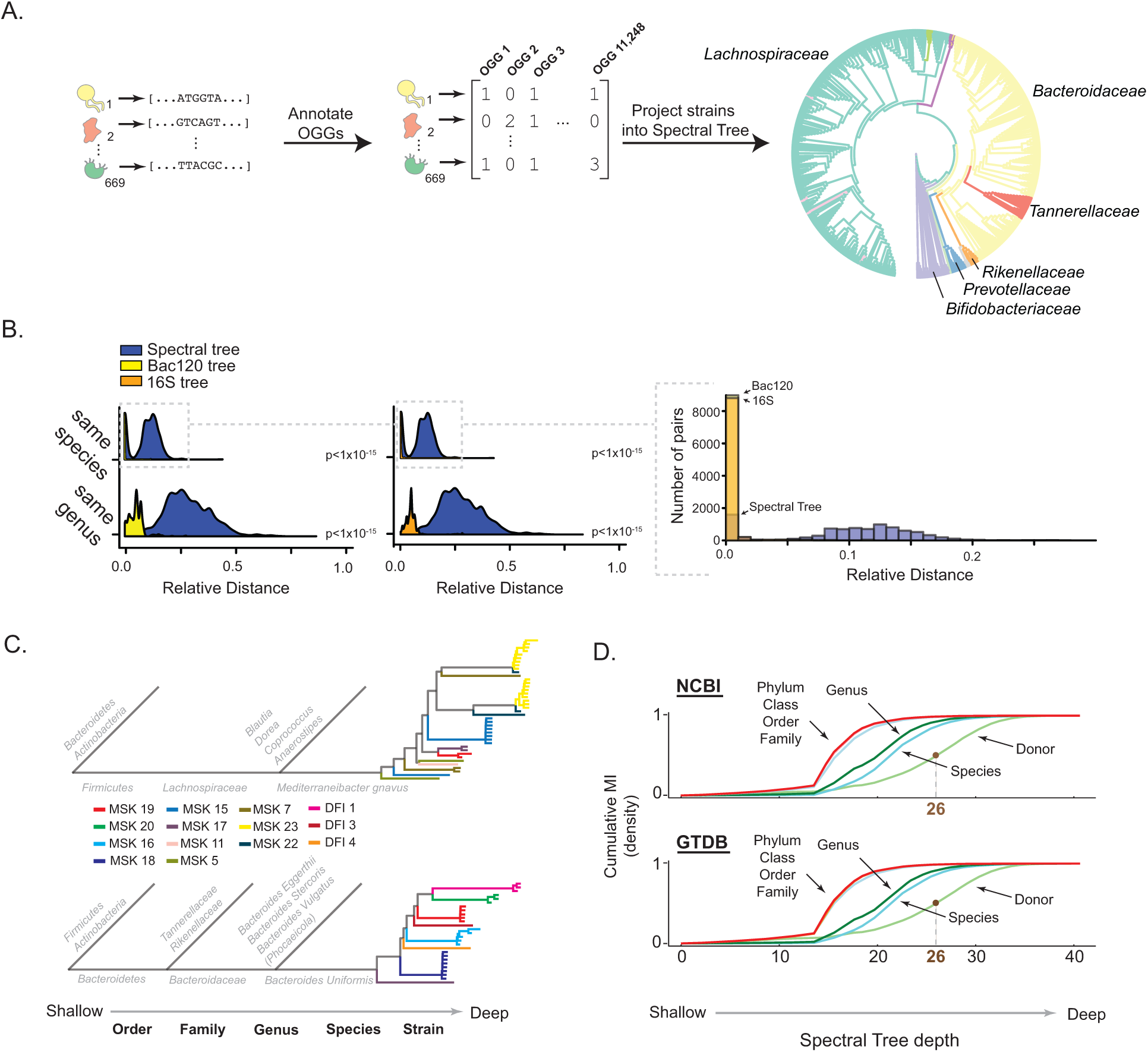
The Spectral Tree reveals subspecies phylogeny. **(A)** Workflow for projecting commensal strain bank into the Spectral Tree. **(B)** Distributions of relative distance between all pairs of strains in commensal strain bank belonging to either the same genus (bottom panel) or species (top panel). Relative distance is defined by either (i) the 16S phylogenetic tree (orange distribution), (ii) the bac120 phylogenetic tree (yellow distribution), or (iii) the Spectral Tree (blue distribution). Inset: distribution of relative distances for all strain pairs that are of the same species. **(C)** Following strains of *M. gnavus* (upper panel) and *B. uniformis* (lower panel) from shallow to deep branches of the Spectral Tree. Each leaf is a strain colored by the identity of the donor from which the strain was collected (see color key. MSK indicates donor from Memorial Sloan Kettering Hospital; DFI indicates donor from Duchossois Family Institute). **(D)** Information shared between phylogenetic designation (NCBI or GTDB database) or donor origin (color-key) and depth of strain cluster in Spectral Tree (x-axis). Tree depth at which 50% of cumulative information regarding shared donor identity is represented is delineated (brown).

We next sought to interrogate the structure of strain-level variation within the Spectral Tree. Focusing on the group of 41 *M. gnavus* strains in our strain bank, we found that the Spectral Tree defined phylogenetic structure through species-level designation, but also showed statistically significant non-random clustering amongst strain-level variants. Interestingly, we found a direct relationship between strain-level variants and donors from which strains were collected (**Fig. 2C**, upper panel). In another example, the 27 strains of *Bacteroides uniformis* illustrated the same trend of being clustered by donor origin (**Fig. 2C**, lower panel). To test the generality of this result across the commensal strain bank, we performed a systematic analysis by computing the Mutual Information (MI) between strain clusters from shallow to deep in the Spectral Tree and whether the clusters shared the same phylogenetic designation or donor origin (Methods). We found that the pattern of strain clustering across the tree reflected a distinct biological order: shallow clusters reflected broad phylogenetic differences, deeper clusters reflected finer phylogenetic differences, and the deepest clusters reflected variation between strains of the same species but isolated and cultured from different donors (**Fig. 2D**). Collectively, these results showed that the deepest Spectral Tree clusters revealed subspecies phylogeny in our strain bank and that subspecies phylogeny was driven by the econiche of individual hosts.

In totality the Spectral Tree contained 41 layers. The layer at which subspecies phylogeny was defined was layer 26 (**Fig. 2D**). As the Spectral Tree was built from >7,000 principal components spanning over 10,000 OGGs, we sought to understand what properties underlie the substantial compression of information we observe. To delineate the pattern of OGGs that define hierarchical relationships in the Spectral Tree, we identified OGGs that were significantly differentially abundant between daughter branches of a given cluster (**Fig. S12**). Interrogating the pattern of OGGs across clusters in the Spectral Tree, we found that the Spectral Tree is organized is through nested genomic variation. That is, variation in OGGs defining the second layer of the Spectral Tree is nested within OGGs defining a cluster in the first layer. This hierarchical pattern continues until the last layer of the tree (**Fig. S13A**) (Methods). Crucially, this property of nestedness enabled explicitly identifying genomic differences that distinguish clusters of strains—a property we used to examine subspecies phylogeny as described next (**Fig. S13B**).

### Functional and evolutionary characterization of subspecies phylogeny

What are the origins of structured phylogeny below the level of species? We used the Spectral Tree to better understand drivers of subspecies phylogeny within our strain bank. As an example, our strain bank contained 20 strains of *Eubacterium rectale* collected from several donors. We isolated the Spectral Tree branch that separated different groups of *E. rectale* strains. As expected per our results in **Fig. 2D**, the groups of strains clustered by donors from which they were isolated (MSK17 and MSK22 versus MSK16, MSK13, and MSK9; ‘MSK’ stands for Memorial Sloan Kettering, the hospital from which donors were recruited and fecal samples were isolated) (**Fig. 3A**). Differences in OGGs between the strains of *E. rectale* illustrated a pattern of mutually exclusive presence or absence. Strains isolated from donors MSK22 and MSK17 harbored gene groups associated with directed motility, with many gene groups encoding structural elements of the flagellum, chemotaxis machinery, and associated signaling cascades. In contrast, strains derived from MSK13 and MSK9 lacked many gene groups encoding components of motility and instead contained gene groups associated with the presence of phage—phage plasmid primase activity, DNA methyltransferase activity, and type I restriction modification. Strains from MSK16 were unique; these strains harbored a subset of gene groups associated with motility but also several gene groups associated with the presence of phage. Collectively, the pattern of gene group presence/absence distinguished *E. rectale* strains hierarchically. Strains from MSK22 and MSK17 were more like each other than strains from MSK16, MSK13, and MSK9; strains from MSK13 and MSK9 were more similar than strains from MSK16.

**Fig. 3.**
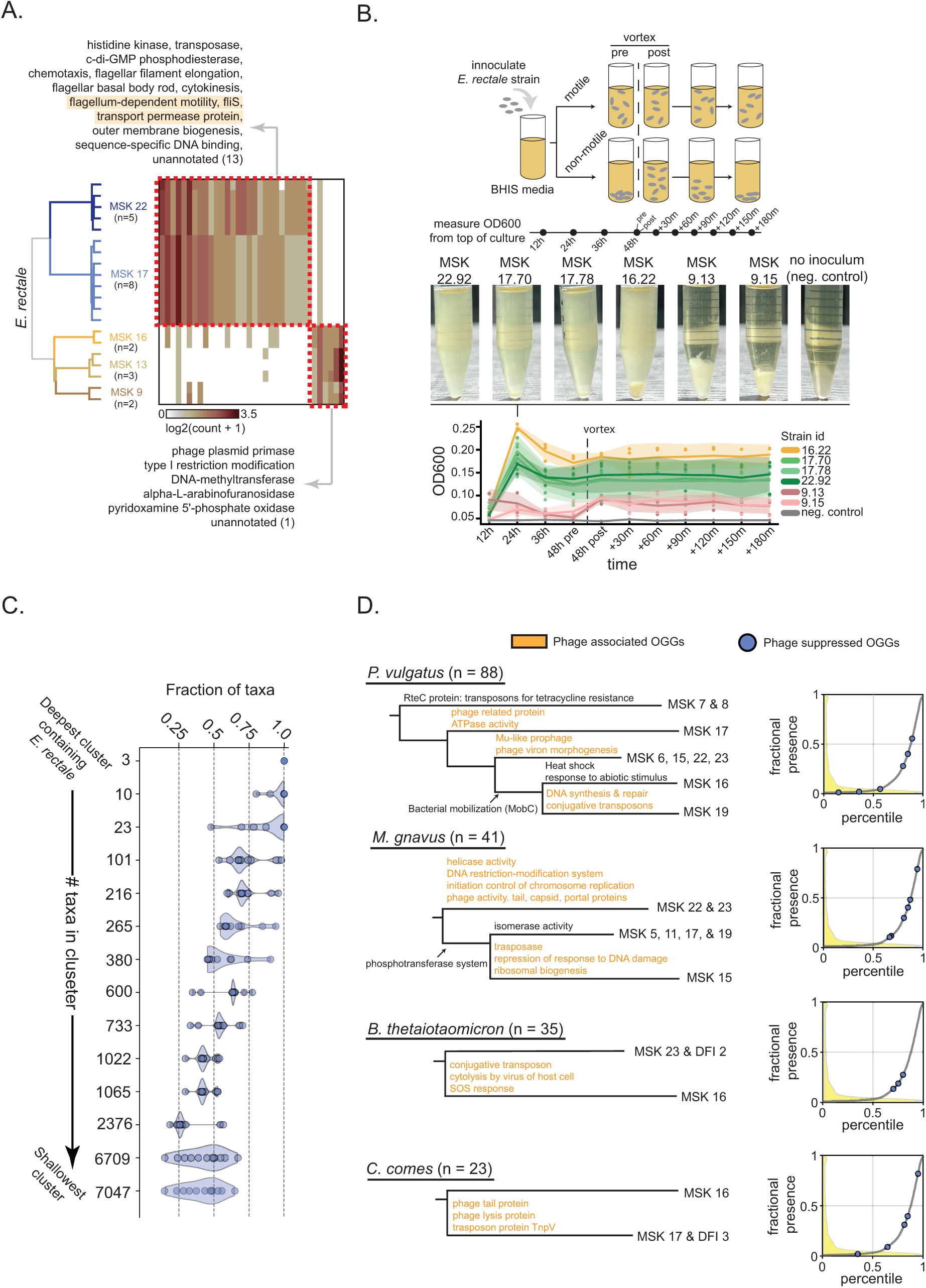
Functional and evolutionary characterization of subspecies phylogeny. **(A)** Clusters of *E. rectale* strains from the Spectral Tree. Branches are colored by strain cluster and are labeled by the donor from which they were isolated. Number in parenthesis below each donor is number of strains. Heatmap shows gene groups that are significantly differentially abundant between strains. Functional annotations of gene groups defining each cluster (red boxes) are shown in text. Highlighted annotations reflect gene groups shared amongst strains from MSK22, MSK17, and MSK16. **(B)** Evaluating motility of *E. rectale* strains derived from different donors. BHI media is inoculated with strains, grown for 48 hours, vortexed, then observed for 180 minutes. OD_600_ measurements are taken from the top of the culture. Pictures show cultures of six different strains—three from MSK22 and MSK17, three from MSK16 and MSK9—and a negative control of media alone after being grown for 24 hours. OD_600_ (y-axis) versus time for each strain in triplicate is shown. Solid lines are average OD_600_ value, contours are ± 1 standard deviation from average OD_600_ value. **(C)** The fraction of taxa (x-axis) containing the 12 annotated OGGs (circles) absent in MSK16, MSK13, and MSK9 out of all taxa within a given cluster in the Spectral Tree (y-axis). y-axis is ordered from the deepest cluster containing the reference *E. rectale* proteome (top) to the shallowest cluster (bottom). **(D)** Left panel; Spectral Tree for given species. Leaves are labeled by donors from which strains were collected; number of strains collected for each species indicated in parenthesis. Text along branches indicate functional annotation of significantly differentially abundant OGGs between daughter clusters. Orange text indicates annotations associated with phage presence; black text along daughter cluster indicates functional annotations of OGGs that are absent termed ‘Phage suppressed OGGs’. Right panel; all 10,117 OGGs are ordered by their percentile rank of fractional presence in the UniProt database (x-axis) and plotted against their fractional presence (y-axis) (grey distribution). The density of OGGs for a particular percentile rank is shown in the yellow distribution. Phage suppressed OGGs for each species are plotted along the grey distribution in blue circles.

The statistically deduced patterns of gene group presence/absence motivated testing *E. rectale* isolates from these donors for their motility. Isolates were tested for their ability to swim in liquid Brain Heart Infused (BHIS) media (Methods). Six strains, three from each major cluster in **Fig. 3A**, were grown anaerobically for 48 hours, vortexed, then observed for 180 minutes (**Fig. 3B**, top panel). After 24 hours of growth, cultures inoculated from strains of MSK22 and MSK17 were uniformly turbid illustrating the robust motility of *E. rectale* strains, while those inoculated with strains from MSK9 exhibited a large pellet with clear inoculum. The culture inoculated with strains derived from MSK16 exhibited a phenotype following their pattern of OGG presence/absence—pellet formation with uniform turbidity—illustrating the compromised ability to swim likely due to the absence of the basal body and other key flagellar and motility components (**Fig. 3B**, middle panel). The OD_600_ measurements of each culture after vortex were in accord with the phenotypes expected from the pattern of gene group presence/absence for each strain (**Fig. 3B**, lower panel). Findings in liquid media were also consistent with motility tests performed in solid agar (**Fig. S14**). These results demonstrated that the subspecies phylogeny amongst *E. rectale* strains manifest as biological differences that could be directly interpreted and learned from the Spectral Tree.

A previously published analysis of *E. rectale* strains demonstrated that a majority of the clade contained motility genes, excepting a single European subspecies (*33*). Thus, our result suggested that subspecies phylogeny associated with phage infection may correlate with strain differences in well conserved areas of bacterial genomes. We examined the conservation pattern of the 12 annotated gene groups that were absent in *E. rectale* strains isolated from donors MSK16, MSK13, and MSK9 but present in strains isolated from donors MSK22 and MSK17 across the entire Spectral Tree. We found that in the phylogenetic local vicinity of *E. rectale*, the 12 gene groups were well-conserved, found in 100% of strains. As we expanded from this vicinity and progressively included more phylogenetically distant bacteria, we found that the 12 gene groups maintained their high conservation, spanning a fractional presence of 20% to greater than 50% across all 7,047 bacteria within UniProt (**Fig. 3C**). These results highlighted that phage-related differences amongst strains associated with variation amongst highly conserved *E. rectale* genes.

We then performed a more systematic analysis, focusing on five species outside of *E. rectale* that were represented by more than 20 strain-level variants where differences amonst gene groups were significant with respect to effect size (log-fold-change greater than 1). These species were *B. uniformis*, *Phocaeicola vulgatus*, *M. gnavus*, *Bacteroides thetaiotamicron*, and *Coproccocus comes*, comprising 214 strains in total. We found that the most conserved gene groups defining subspecies phylogeny for all species were related to phage physiology. Other features included gene groups related to horizontal gene-transfer and inter-cellular competition, amongst many other annotations (**Table S3**). These results were consistent with previous metagenomic-based analyses of subspecies variation in human gut microbiomes illustrating the importance of phage in mediating strain-level variation (*12*). We also found that the presence of phage elements correlated with the absence of gene groups that are phylogenetically conserved. We term ‘Phage-suppressed’ OGGs as gene groups whose absence was shared with the presence of phage-related gene groups. Across all species that were analyzed, the Phage-suppressed OGGs were predominantly within the top half of gene groups ranked by fractional abundance across all 10,177 OGGs defining bacterial proteomes in UniProt. Additionally, several Phage-suppressed groups were present in greater than 20% and up to 80% of all taxa in UniProt (**Fig. 3D**, right panel), illustrating their broad conservation across the Kingdom Bacteria.

Collectively, these results suggest that subspecies phylogeny is markedly associated with a shared history of phage exposure amongst groups of donors and manifests as functionally relevant changes in clusters of strains. Our findings illustrate an environmental origin for how strain-level variation becomes structured below the phylogenetic level of Species.

### Predicting the metabolic capacities of strains

We tested whether the Spectral Tree could be used to relate genotypes of individual strains with their metabolism. To address this idea, we first constrained our analysis to species where we had at least 20 representative strains in our strain bank for statistical power. In total, this amounted to 356 strains across 11 species. We then labeled all 356 strains according to their branching pattern in the Spectral Tree. Since strains are linked in the Spectral Tree by a common root, the branching pattern of each strain is a unique ‘barcode’ of statistically inferred lineage (**Fig. 4A**). We therefore termed this barcode a ‘Spectral Lineage Encoding’ (SLE). We then trained LASSO models that used the SLE for each bacterial strain as input and the relative difference in metabolite concentrations for all metabolic features that we profiled as output (**Fig. 4B**) (Methods). To ensure an out-of-sample prediction for all strains, we computed the spectral distance of a randomly chosen 75% of the 356 strains within the Spectral Tree as a training set. Next, we assigned SLEs to all strains in the training set, trained a LASSO model, and validated the LASSO model on the remaining 25% of strains. We then repeated these steps four times across five different re-partitions of the dataset (**Fig. S15**) (Methods). For comparison, we also measured the predictive capacity of LASSO models trained on either the top three or top 10 principal components of strains within our strain bank annotated by their OGG content (Methods).

**Fig. 4.**
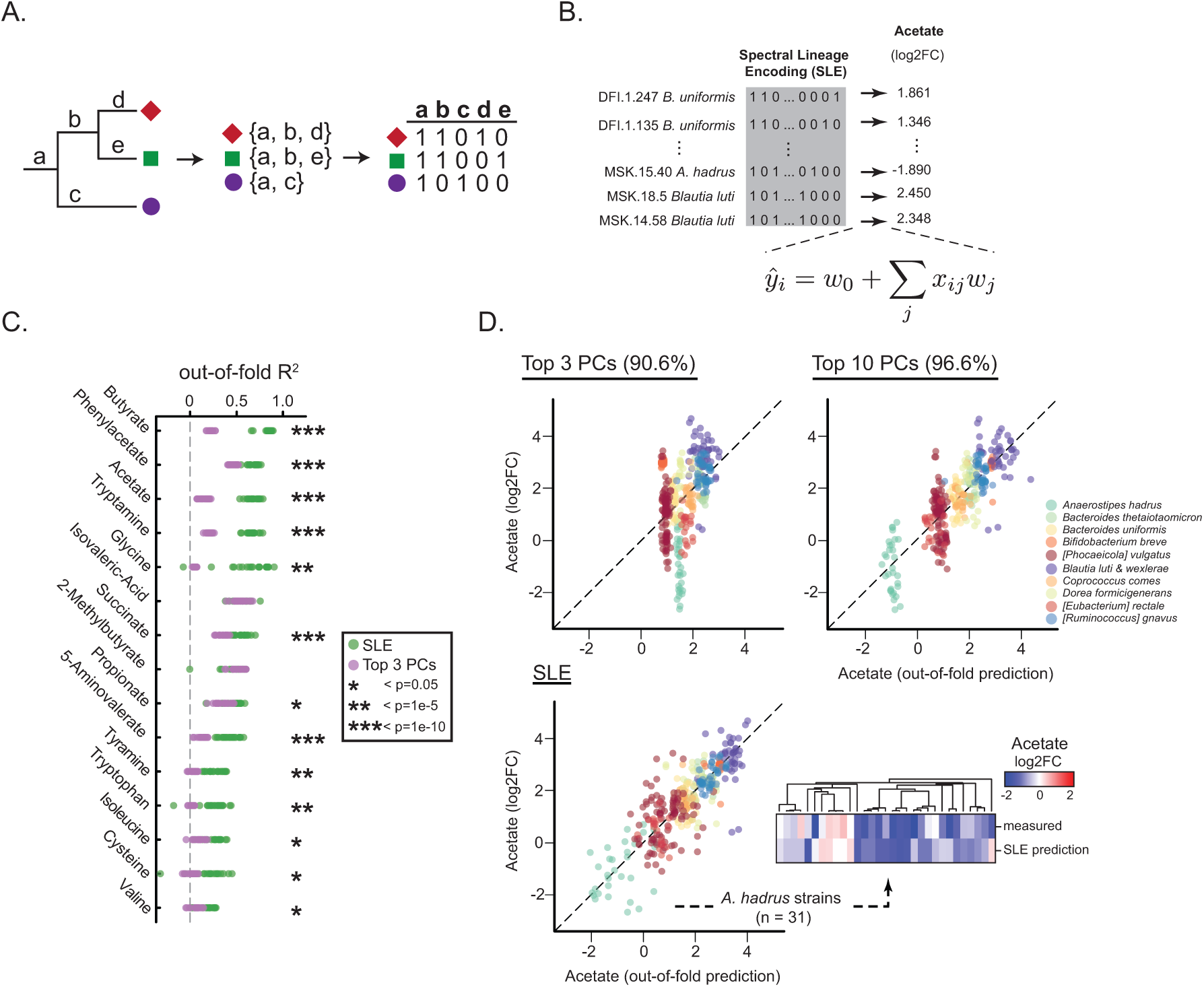
Predicting metabolic traits of individual strains from the Spectral Tree. **(A)** Workflow for defining Spectral Lineage Encodings (SLEs) for taxa. Taxa (red diamond, green square, purple circle) are labeled according to their unique branching path along the Spectral Tree and a table of SLEs is created with branches as features comprising the columns and taxa as rows. **(B)** Schematic for training a LASSO model on SLEs to predict acetate fold-change (FC) for strains. **(C)** Predictive capacity (out-of-fold R^2^, x-axis) of 20 SLE LASSO models (green circles) for 15 metabolites (y-axis) and 20 LASSO models trained on top three principal components (PCs) of strain co-evolution (purple circles). * indicates degree of statistically significant difference between predictive capacities of models (see key). **(D)** Out-of-fold prediction (x-axis) versus measured fold-change (y-axis) for acetate metabolism across all strains (see color key for species designation) for LASSO models trained on top three principal components (harboring 91% variance), top 10 principal components (harboring 97% variance), and SLEs. Inset shows predicted and measured fold-change of acetate for 31 *Anaerostipes hadrus* strains; strains are hierarchically clustered by fold-change measurement.

Our results showed that for several metabolic features, the LASSO models trained on SLEs did not predict relative metabolite consumption or production over an r^2^ value of 0.5 (**Table S4**). However, for 13 metabolites spanning short-chained fatty acids, amino acids, and indoles, the LASSO models on SLEs accurately predicted strain-level metabolic output (**Fig. 4C**). Using relative concentrations of acetate as an example, we found that LASSO models trained on the top three principal components collapsed predictions according to coarse phylogenetic trends. The group of LASSO models trained on the top 10 principal components collapsed predictions according to more resolved phylogenetic trends. In contrast, the group of SLE LASSO models resolved strain-level metabolic variability across the full dynamic range of fold-change (**Fig. 4D**). As an example, 31 strains of *Anaerostipes hadrus* exhibited strain-level variability in acetate metabolism with many strains consuming acetate while a minority of strains produced acetate. Comparing the predicted with observed relative concentrations of acetate illustrated that the LASSO model trained on SLEs correctly predicted the cluster of *A. hadrus* strains deviating from the coarse phylogenetic trend (**Fig. 4D**, inset). Collectively, these results showed that describing strains by their co-evolutionary signature defined by the Spectral Tree— the SLE—resolves strain-level metabolic variability.

### Using SLEs to rationally engineer consortia

Our findings framed an approach for predicting the metabolic capacity of newly acquired bacterial strains from their genome content alone. First, sequence the newly acquired strain and annotate by OGGs; second, project into the Spectral Tree; third, compute SLE; and fourth use the SLE as an input to our LASSO model. One rationale for creating gut bacterial strain banks has been to design consortia for specific purposes. We reasoned that the SLE-based LASSO models could enable rationally engineering communities of strains for desired metabolic properties. That is, from a strain bank of sequenced strains, our approach could inform which strains to add to a consortium to yield a desirable metabolic output from genome sequence alone.

As a proof-of-concept of our idea, we constructed 17 different consortia comprised of three bacteria each. Each consortium contained a uniform background of two strains— *Clostridium scindens* plus *Bifidobacterium longum*—and one of 17 possible phylogenetically diverse strains that we isolated and cultured from human fecal samples (**Fig. 5A**, left panel) (Methods). We whole genome sequenced each of the 17 strains and annotated their OGG content. Eight of the 17 strains were related to at least one representative member of the same genus in our commensal strain bank while 9 strains were from genera outside our strain bank (**Fig. S16**). The metabolic output for each of the 17 strains was predicted using our SLE LASSO models. We compared the metabolic output of the bacterial consortium with the predicted metabolic output of the third strain (**Fig. 5A**, right panel). We focused our analysis on acetate concentrations as our SLE LASSO models showed the best predictive capacity at the strain-level for this metabolite. *C. scindens* and *B. longum* each produced acetate individually and increased acetate production in combination (**Fig. S17**), **Table S5**].

**Fig. 5.**
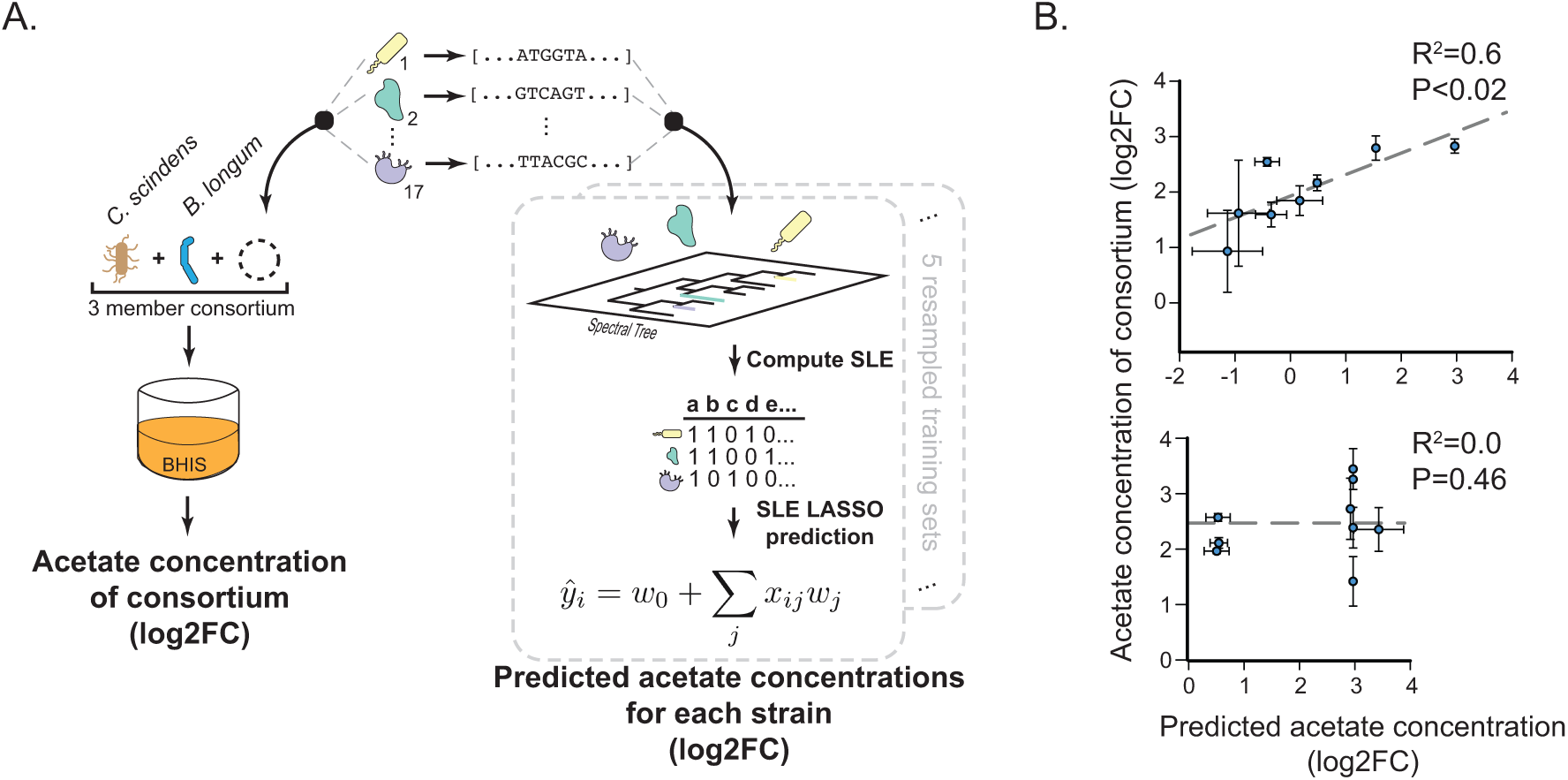
SLEs enable rationally engineering consortia from strain genomes. **(A)** 17 new bacterial strains were isolated from fecal samples, whole genome sequenced, and added to a consortium of *Clostridium scindens* and *Bifidobacterium longum*. Consortia were grown and acetate concentrations were measured (left panel). Acetate concentrations for all 17 strains were predicted using SLE LASSO models (right panel). **(B)** Predicted relative acetate concentration for each of the 17 strains (x-axis) versus measured relative acetate concentration for cultured consortia (y-axis). Strains are grouped by whether the commensal strain bank contained a strain with a related genus (upper panel) or did not contain any strain with a related genus (lower panel).

Our results fell into two separate classes depending on whether the third strain was related to other strains within our strain bank. For strains where there was an existing strain in the same genera, we found that the acetate concentrations of the consortium were predictable from the predicted acetate output of the third strain (**Fig. 5B**, upper panel). In contrast, for strains that were unrelated to any strain in the strain bank, we found no correlation between predicted strain-level acetate concentrations and consortium acetate concentrations (**Fig. 5B**, lower panel). Our findings suggest that our genome-based models were informative for rationally engineering small bacterial communities with predictable metabolic traits, suggesting the possibility of ‘bottom-up’ consortia design—building up complex communities using individual strains—from genome content alone (*34*, *35*).

Our data suggest that given a model that is sufficiently predictive of strain phenotype, considering higher-order microbial interactions may not be necessary for microbial community design. This observation is consistent with recent work showing that phenotypes of microbial communities can predicted and learned from considering just first-order and pairwise interaction terms (*36*, *37*). In addition, the utility of our models does not generalize beyond strains that are not phylogenetically represented in our strain bank. This finding suggests that collecting, sequencing, and phenotyping a broad phylogeny of strains, as opposed to focusing data collection efforts on increasing coverage of strains within a given species, may be a useful strategy to build statistical models for engineering complex consortia.

## Discussion

The importance of individual strains in mediating gut microbiome function requires new frameworks for their description beyond merely taxonomic definitions. Here, we showed that co-evolutionary patterns derived from a large diversity of strains across the bacterial tree-of-life creates a natural, data-driven, and useful description of gut commensal strains. Importantly, our findings demonstrate how leveraging biological diversity reflective of many diverse and unrelated environments can help understand constraints on genomic variation within the single environment of the human gut. As our framework is not specific to gut bacteria but can be applied to strains isolated from any environment, we pose that the construct we have developed—the Spectral Lineage Encoding—may be a generally useful schema for describing and studying bacterial strains.

The intra- and interpersonal variation in the structure of human gut microbiomes has been extensively described (*38–40*). The degree to which this variation reproducibly derives from external factors has remained a subject of discussion with recent studies attempting to control for environment—like diet or spatial geography—to ‘normalize’ structural changes observed in human cohort studies (*41–43*). Our data suggest that a history of phage infection can lead to structured, non-random microbiome changes between groups of humans that manifest in subspecies phylogeny. While we demonstrated how these changes lead to different behaviors at the scale of individual bacteria, the functional consequences of such strain-level variation at the scale of the whole microbiome remains to be characterized. Interestingly, though a majority of the Phage-suppressed OGGs we identified were phylogenetically conserved, the strains nevertheless persisted in the gut microbiome of donors. This suggests that perhaps changes in conserved genomic areas within individual bacteria can be tolerated without a substantial fitness decrease when considered within the context of the entire gut ecosystem. The recent shift towards genomic analysis of bacteria within the context of whole microbial ecosystems will enable a better definition for a ‘null hypothesis’ of genomic constraint within individual strains.

What fundamental properties underlie the utility of describing strains by their co-evolutionary signature? Unlike engineered systems, existing or ‘extant’ biological systems arise from ancestors through the evolutionary process (*44–48*). Therefore, understanding how patterns of genetic interactions encode behaviors is inextricably intertwined with defining commonalities and divergences in molecular structure (*32*, *49–54*). Current statistical strategies for parsing differences amongst genomes involve so-called ‘factorization’ approaches that discover low-dimensional representations of high-dimensional patterns of variation. Indeed, the era of biological big-data has seen an explosion in the use of factorization methods (*55*). Such approaches are predicated on a key assumption: the systems being interrogated are unrelated to each other. For evolved systems, ancestral relatedness violates the assumption of system independence, demanding a new formalism for comparative efforts. We reason that the SLE is a useful descriptor of bacteria because it embeds hierarchical scales of relatedness across the evolutionary record, simultaneously capturing both broad phylogenetic differences and fine-grained differences within species. The hierarchical nature of the SLE as a descriptor therefore distinguishes variation arising from ancient phylogenetic sources from functional differences reflecting recent adaptations (i.e. to individual human hosts). This capacity to separate sources of variance is key for creating accurate predictive models of biological behavior from genome content (*25*). We anticipate that our approach is unlikely to be the only applicable framework given the recent development, implementation, and success of large language models (LLMs) in characterizing evolutionary relationships among complex biological systems (*56*, *57*). However, our findings show that creating statistical representations of the evolutionary record may lay a foundation for understanding and predicting idiosyncrasies of individual biological systems that deviate from broad phylogenetic trends. Future studies applying the concepts developed here to other evolved systems will test this idea.

## Acknowledgments

We thank S. Kuehn, M. Mani, D. Pincus, A. Murugan, J. Gordon, S. Oakes, A. Drummond, O. Rivoire, and R. Ranganathan for helpful discussions.

## Author contributions

B.D. and A.S.R. designed this study and conceived of the approach taken. B.D. and R.Y.C. conceptualized ‘spectral distance’ as a quantitative metric. H.G. and B.D. performed motility experiments of *E. rectale* strains. B.B., A.S., and H.L, oversaw the collection, isolation, sequencing, and bioinformatic analysis of the CSB. V.B. conceptualized and wrote the code for evaluating statistically significant differences in feature abundances between daughters of Spectral Tree. A.S. aided in execution and supervision of metabolomic profiling on strains. E.G.P. oversaw and supervised collection, isolation, sequencing, and metabolomic profiling of strains. B.D. wrote all code and conducted all analysis. B.D. and A.S.R. wrote the paper.

## Data and materials availability

All sequencing data can found and downloaded on NCBI. All raw data from metabolomics can be downloaded from Metabolights (Study ID: MTBLS7771). All code used in this study was made in Julia. All code used for genome sequencing and metabolomic processing was made in Python. All code, along with annotations and step-wise instructions, are available for download via a github repository: https://github.com/aramanlab/Doran_etal_2022. Links for specific datasets in Dryad can be found within the same github repository.

