## Supplemental Information for "An evolution-based framework for describing human gut bacteria"

### Materials and Methods

Creating a bank of commensal human gut microbiome strains

Fecal samples were obtained from 28 human donors that fell within the age range of 18 to 63 with a median age of 35. Donors were selected as those with no antibiotic use in the past year, no known history of diabetes, colitis, autoimmune disease, cancer, pneumonia, dysentery, or cellulitis at time of consent. Institutions that approved protocols of fecal sample collection were Memorial Sloan Kettering (MSK) and the University of Chicago. Fresh fecal samples were immediately reduced in an anaerobic chamber upon collection and diluted and cultured on various growth media. Agar media types vary, but include any of following: Columbia Blood Agar, Brain Heart Infusion +Yeast, Brain Heart Infusion + Mucin, Brain Heart Infusion + Yeast + Acetate or N-Acetylglucosamine, reinforced Clostridial Agar, Peptone Yeast Glucose, Yeast Casitone Fatty Acids, Defined media M5. Colonies were selected and grown to be sufficiently turbid, 20% glycerol/PBS stocks were created and stored in a -80°C freezer.

Colonies were selected for whole-genome based on pyro-sequencing of the 16S region which provides a rough estimate of genus level designation. For each donor, only colonies that had a sequence identity threshold of less than 99% from CD-Hit (v. 4.8.1) were selected for whole-genome sequencing (*1*). Bacterial genomic DNA was extracted using QIAamp DNA Mini Kit (QIAGEN) according to manufacturer's manual. The purified DNA was quantified using a Qubit 2.0 fluorometer. 1000ng of each sample was prepared for sequencing using the QIAseq FX DNA Library Kit (QIAGEN). The protocol was carried out for a targeted fragment size of 550bp. Sequencing was performed on the MiSeq or NextSeq platform (Illumina) with a paired-end (PE) kit in pools designed to provide 1-3 million PE reads per sample with read length of 250 or 150 bp. Adapters were trimmed off with Trimmomatic with following parameters: the leading and trailing 3 bp of the sequences were trimmed off, quality was controlled by a sliding window of 4, with an average quality score of 15 (default parameters of Trimmomatic). Moreover, any read that was less than 50 bp long after trimming and quality control were discarded. The remaining high-quality reads were assembled into contigs using SPAdes (v3.14.0)(*2*).

Taxonomic classification of the assembled contigs was performed with the following methods: (a) Kraken2 (v2.1.1); (b) full/partial length 16S rRNA gene from each isolated colony's assembled contigs is extracted and input into BLASTn (v2.10.1+) to query against NCBI's RNA RefSeq database (*3*, *4*). Top five hits for each query are manually curated to determine an isolate's identity, with identity and coverage cutoff both at 95%; (c) GTDB-Tk (v1.5.1) (*5*). Final taxonomy is determined by the consensus of the three methods. Any colony that did not match initial pyro-sequencing taxonomy or lacked consensus are excluded from the commensal strain bank.

Metabolic profiling of the commensal strain bank

Strains were grown until sufficiently turbid then spun down. Supernatant samples were frozen at -80 °C prior to extraction. Samples were thawed and extraction solvent (4 volumes of 100% methanol spiked with internal standards and stored at -80 °C) was added to the liquid sample (1 volume) in a microcentrifuge tube. Tubes were then centrifuged at -10 °C, 20,000 x g for 15 min and supernatant was used for subsequent metabolomic analysis. Compounds were derivatized as described by Haak *et al.* with the following modifications (*6*). The metabolite extract (100 μL) was added to 100 mM borate buffer (100 μL, pH 10), 100 mM pentafluorobenzyl bromide in acetonitrile (400 μL), and n-hexane (400 μL) in a capped mass spectrometry autosampler vial. Samples were heated in a thermomixer C (Eppendorf) to 65 °C for 1 hour while shaking at 1300 rpm. After cooling to room temperature, samples were centrifuged at 4 °C, 2000 x g for 5 min, allowing phase separation. The hexanes phase (100 µL) (top layer) was transferred to an autosampler vial containing a glass insert and the vial was sealed. Another 100 µL of the hexanes phase was diluted with 900 µL of n-hexane in an autosampler vial. Concentrated and dilute samples were analyzed using a GC-MS (Agilent 7890A GC system, Agilent 5975C MS detector) operating in negative chemical ionization mode, using a HP-5MSUI column (30 m x 0.25 mm, 0.25 µm; Agilent Technologies 19091S-433UI), methane as the reagent gas (99.999% pure) and 1 µL split injection (1:10 split ratio). Oven ramp parameters: 1 min hold at 60 °C, 25 °C per min up to 300 °C with a 2.5 min hold at 300 °C. Inlet temperature was 280 °C and transfer line was 310 °C. A 10-point calibration curve was prepared with acetate (100 mM), propionate (25 mM), butyrate (12.5 mM), and succinate (50 mM), with 9 subsequent 2x serial dilutions. Data analysis was performed using MassHunter Quantitative Analysis software (version B.10, Agilent Technologies) and confirmed by comparison to authentic standards. Normalized peak areas were calculated by dividing raw peak areas of targeted analytes by averaged raw peak areas of internal standards. GC-MS was used to detect compounds following pentafluorobenzyl bromide (PFBBr) derivatization. Normalized relative abundance values (50 compounds from SCFA, BCFA, amino acid, aromatic, hydroxylated fatty acid, organic acid, indole, and additional subclasses) are reported following PFB derivatization and detection by negative collision induced-gas chromatography-mass spectrometry ((-)CI-GC-MS, Agilent 8890).

<u>Creating phylogenetic trees of commensal strain bank using 16S and bac120 proteins</u>

The 16S sequence was isolated from each strain in the commensal strain bank. All 16S sequences were aligned with Mafft (v7.520) creating a multiple sequence alignment of 1521 features and 335 unique sequences (*7*). This alignment was then input to phyml (v3.3.20200621) with these command options: *"./phyml -dnt -mHKY85 -fe -o tlr --search SPR --r_seed 123456 --rand_start --n_rand_starts 3 --no_memory_check --bootstrap -4 -i BB669_16S.phy"* (*8*). Redundant sequences were placed into the final tree by PhyML at distance zero from their identical representative in the tree.

For bac120, the fastafiles for each isolate was input to the gtdbtk (v2.3.0) identify and align pipeline to create a multiple sequence alignment with the bac120 feature set comprising 5,035 features, with 311 unique sequences (*9*). This alignment was input to PhyML (v3.3.20200621) with these command parameters: *"./phyml -daa -mLG -fe -o tlr --search SPR --r_seed 123456 --rand_start --n_rand_starts 3 --no_memory_check --bootstrap -4 -i BB669_bac120.phy"* (*8*). Redundant sequences were placed into the final tree by PhyML at distance zero from their identical representative in the tree.

<u>Annotating each strain in the commensal strain bank by its orthologous gene group (OGG) content</u>

For individual isolates, the genome assemblies were annotated using Prokka (v1.12) producing a fasta file of all coding regions from the assembled genome translated to the amino-acid protein sequences (*10*). This fasta file was then input to eggNOG-mapper (v2.0.1b) to annotate each protein sequence against the eggNOG database (v5.0) of orthologous gene-groups (OGGs) at the level of *Bacteria* ('@2') (*11*, *12*). Each isolate was then aligned based on this common set of OGG features, where isolates correspond to each row and OGGs correspond to each column and each entry holds the number of protein sequences that matched to that OGG. This OGG alignment of 669 isolates forms the CSB OGG matrix (**Fig. 1C**). Each isolate was annotated across 11,248 total OGGs of which 5,449 OGGs have greater than zero variance; annotations of isolated were done with 16S-BLASTn, GTDB, and final NCBI taxonomic designations at the level of Phylum through Species.

Generating PCA and UMAP plots in **Fig. 1D** and **Fig. S2**

The matrix of commensal strain bank strains defined by their OGG content was progressively subset from the whole strain bank to (i) the family *Bacteroidaceae* (n = 229 strains), (ii) the genus *Phocaeicola* (n = 103 strains), and (iii) the species *P. vulgatus* (n = 93 strains). For each subset, the strains were analyzed using PCA; resulting plots are shown in **Fig. 1D**. For the whole strain bank, PCA and UMAP analysis was performed; resulting plots are shown in **Fig. S2**.

Measuring differences in compute time between methods for defining bacterial phylogenetic relatedness

To evaluate the performance of different methods of phylogenetic tree inference, we constructed 4 alignments of increasing numbers of taxa. These alignments were created by sub-sampling an alignment of 7,047 UniProt reference proteomes annotated by Orthologous Gene Group which was defined in previous work published by our laboratory in Zaydman *et al.* (*13*). We subsampled to (i) 25 taxa from Genus *Ruminococcus*, (ii) 50 taxa from the Family *Rhodospirillaceae*, (iii) 103 taxa from the Order *Oceanospirillales*, and (iv) 211 taxa from the Class *Bacteroidia*. We then transformed the count matrices associated with each of the alignments into an alignment where if the count of an OGG is greater than 0, the account is assigned a thymine ('T') and otherwise the count is assigned an adenine ('A'). This transformation is performed to facilitate use of commonly used phylogenetic tree-building methods. Each MSA is used as input for PhyML v3.3.2, FastTree v2.1.11, RAxML v8.2.12, MrBayes v3.2.7, and our spectral approach with the clock time in seconds recorded for each tool (*8*, *14*–*16*). This analysis was performed in parallel on the University of Chicago Midway2 cluster (2.4 GHz 7 core Intel E5-2680 v4 CPU with 16 GB of RAM per tree building task).

Constructing a Spectral Tree from 7,047 bacterial reference proteomes in UniProt

Construction of the full alignment of 7,047 UniProt reference proteomes annotated by 10,177 Orthologous Gene Groups was previously described in Zaydman *et al*. (*13*). For this alignment we followed the procedure outlined in *Supplementary Text: (Section 1.3) Creating a tree of relatedness from covariation structure of taxa* to create a Spectral Tree.

Projecting commensal strain bank into Spectral Tree of 7,047 UniProt reference bacterial proteomes

Genome sequences of all strains from the commensal strain bank were put through EggNog mapper (emapper 5.0) and their proteomes were annotated for their OGG content across the same set of OGGs defining the UniProt reference proteome database (n = 10,177). Any OGG measured in UniProt but not in the commensal strain bank was imputed as a '0' count. From **eq. (1)** in Supplementary Text, we calculated the principal components defining covariation amongst the UniProt reference proteomes, $P^{UniProt}$, as

$$P^{UniProt} = D^{OGG} \, V^{UniProt} \tag{2}$$

where $D^{OGG}$ is the 7,047 UniProt reference bacterial proteomes annotated by their 10,177 OGGs.
We next defined $B^{OGG}$ as the matrix of commensal strain bank strains annotated by their OGGs. Therefore from (2),

$$P^{CSB} = B^{OGG} \, V^{UniProt} \tag{3}$$

where $P^{CSB}$ is a matrix of commensal strains (rows) by 7,047 principal components (columns) that collectively define the structure of bacterial co-evolution in the UniProt database; each entry is the contribution of each commensal strain bank strain onto each principal component.

Computing the information shared between clusters defined by the Spectral Tree and shared properties of strains within clusters

We first created 100 'cuts' of the tree where each cut is equally spaced across the depth of the tree. The first cut is defined at the root of the tree forming a single cluster comprising all taxa; the last cut is defined at the terminal branches of the tree forming as many clusters as there are taxa. For each cut, we form two membership vectors **C** and **T** where each element in the vector represents a pair of taxa in the tree. **C** is a '1' if two taxa belong to the same tree cluster and a '0' if not. **T** is a '1' if two taxa share a property (i.e. belong to the same NCBI taxonomic designation or come from the same donor) and a '0' if not. **T** is constructed for (i) all taxonomic designations spanning 'Phylum' to 'Species' in NCBI, (ii) all taxonomic designations spanning 'Phylum' to 'Species' in GTDB, and (iii) identity of donor. We then calculate the mutual information (MI) between **C** and **T** by the following equation:

$$MI(\boldsymbol{C},\boldsymbol{T}) = H(\boldsymbol{C}) + H(\boldsymbol{T}) - H([\boldsymbol{C},\boldsymbol{T}]^t) \qquad (4)$$

where $H(x) = -\sum_{b\in 0,1} p(x_b)\log_2\big(p(x_b)\big)$ represents the Shannon entropy; $p(x_b)$ is the proportion of x that is equal to either '0' or '1' respectively in the distributions defined by either **C** or **T**. H($[\boldsymbol{C},\boldsymbol{T}]^t$) is the joint entropy of **C** and **T**. The 'Cumulative MI density' plotted in **Fig. 2D** is defined by adding the MI for each subsequent deeper cut of the tree and dividing by the total sum of MI across all cuts. NCBI phylogenetic strings were mapped to NCBI taxonomy IDs following methods described in Zaydman et al. (*13*).

To measure uncertainty in MI, we bootstrapped the MI calculations. For a given MI between **C** and **T**, pairs of taxa (matched elements of **C** and **T**) are sampled with replacement to the same total number of pairs. As an example, for 10 choose 2 pairs (n=45), 45 pairs are sampled with replacement. This bootstrapping is performed 10 times and the cumulative mean ± 2 standard deviations of MI is plotted as ribbons.

Motility experiment in *E. rectale*

On day 1, BHIS media (250mL dH2O, 9.25g BHI Media, 2.5mL cysteine solution (1g cysteine in 10mL dH2O)) was made and aliquoted into 50 mL conical tubes. Tubes were cycled into anaerobic chamber (Coy) 24 hours prior to the experiment. Caps on tubes were left loose to allow for equilibration to the anaerobic environment and to release excess oxygen that may impact strain growth. On day 2, a serological pipet was used to aliquot 5 mL BHIS into 20 mL conical tubes. Glycerol stocks of *E. rectale* strains were cycled into the anerobic chamber; media was inoculated with strains in triplicate and placed in a 37° incubator in the anaerobic chamber. $OD_{600}$ was measured every 12 hours for 48 hours for each tube of inoculated media by sampling from the top of the culture (taking 100µL from the top 1mL of the 5mL cultures). During this time, each tube was also observed for pellet formation. After 48 hours of incubation, cultures were briefly vortexed to disseminate any pellet formed at the bottom of the tube. Samples were then collected from the top of each culture (100µL from the top 1mL) and measured for their $OD_{600}$ every thirty minutes for 180 minutes after vortex.

Training SLE LASSO models for predicting the relative concentration of a metabolite

To establish a training and validation set, we selected all strains belonging to the 10 species with 20 or greater biological replicates (n=356), and then further subset to 75% of those strains maintaining relative proportions of species groups (n = 267) with the remaining 25% (n = 89) used as a validation set.

To train the LASSO model, we first generated a Spectral Tree from the training set and created an associated SLE matrix ($\mathbf{SLE_{train}}$, 267 rows by 266 columns) per the diagram in **Fig. 4A**. Each strain was labeled by the fold-change (log2FC) of a specific metabolite. We next estimated the linear coefficients relating SLEs with relative change in metabolite concentration by

$$\hat{w} = \underset{w}{\operatorname{argmin}}(\ |\boldsymbol{SLE_{train}}\, w\ -\ y|_2\ +\ \lambda|w|_1) \tag{5}$$

where $\hat{w}$ is the estimated coefficients, $y$ is the log2FC of a metabolite, and $\lambda$ is the regularization parameter set to $10^{-3}$. We then made predictions for the validation set by finding their nearest neighbor in the training set using spectral distance. The SLE of this nearest neighbor was then used to make a prediction by

$$\hat{y} = \boldsymbol{SLE_{train}}\ \hat{w} \tag{6}$$

**Eq. (6)** represents the out-of-fold predictions for a single fold. We repeat creating a training set, creating an associated Spectral Tree, creating an associated $\mathbf{SLE_{train}}$, and making out-of-fold prediction across 20 re-samplings comprising 4-folds per data partition across 5 re-partitions. This validation procedure guarantees 5 out-of-fold predictions per taxa. We measure the performance of the model using the Pearson correlation ($r^2$) for each re-sampling.

<u>Training LASSO model on top 3 and 10 principal components for predicting relative metabolite concentration</u>

The procedure for creating a training set, training LASSO models, and creating a validation set was the same as described in '*Methods: Training SLE LASSO models for predicting the relative concentration of a metabolite*'. PCA was performed on the alignment of taxa comprising the training set (267 taxa by 5,499 OGGs), LASSO models were trained on taxa projections onto either the top three or top 10 principal components. For the validation set, taxa were projected into either the top three or top 10 principal components of the training set and their corresponding coordinates were used as input into the LASSO model to make out-of-fold predictions.

<u>Construction and metabolic profiling of consortia described in **Fig. 5**</u>

Across all consortia two members are kept constant, *C. scindens* and *B. longum*. One of 17 different strains was included as the third member. Validated freezer stocks of relevant strains from the consortium were isolated on reduced Columbia Blood agar plates in the anaerobic chamber. At 48 hours, once colonies develop, 3-5 well isolated colonies from each strain were resuspended into 300µL reduced phosphate buffered saline. Tubes were vortexed well to ensure there are no clumps. Optical density of the inoculum was adjusted to OD600 0.3-0.5. Once normalized, 100µl of each prepared inoculum (at 1:1:1 volume) was mixed into another tube and this was used as the final inoculum for consortium experiments.

Metabolomic profiling was performed following the preparation of a 96-well plate, where 200 µL of media (Brain heart infusion medium) and 10 µL of prepared inoculum was added into each well. Taurocholaten(TCA) or Taurochenodeoxycholate (TCDCA) at a final concentration of 10μg/ml was spiked into the media to measure bile acids in addition to short chain fatty acids. Controls included media spiked with either bile acid (TCA and TCDCA) for baseline measurements. Plates were sealed with microplate plate sealing adhesive films and run on a BioTek BTLP600 plate reader for 12-24 hours, until stationary phase was attained. At this point plates were removed from the plate reader and spun down using a tabletop centrifuge at 4,300 x g for 5 minutes. To a new plate, 150 μL of supernatant was transferred and assayed for Acetate concentration (mM) as described in '*Methods: Metabolic profiling of the commensal strain bank*'. Log2 foldchange was computed between the acetate concentration (mM) in supernatant of the consortia.

### Supplementary Text

The Supplementary Text contains four sections for describing our approach to developing a quantitative definition of evolutionary distance from patterns of co-evolution and inferring trees of taxonomic relatedness (termed ‘Spectral Trees’ in the main text). Section 1 explores the relationship between the trajectory of sequential diversifications that give rise to taxa and patterns of covariation between taxa. In Section 1, we use the trajectory described in **Fig. S3** as a case study. Section 2 uses a toy model described in **Fig. S4** to provide a mathematical basis for why the structure of covariation amongst taxa is directly related to a hierarchy of relatedness between taxa, leading to the generality of our findings across diverse taxonomic alignments as described in **Fig. S5**. Section 3 demonstrates how our statistical approach of inferring relatedness between taxa resolves complex diversification trajectories including convergent processes by using the trajectory described in **Fig. S6** as a model. Section 4 provides a stepwise workflow for computing a Spectral Tree from an alignment of biological systems. We used this workflow to create the Spectral Tree comprising 7,047 non-redundant reference bacterial strains from the UniProt database.

#### 1. Relating diversification histories of taxa with patterns of covariation between taxa.

Previous work from our laboratory indicated a relationship between the principal component spectrum of bacterial co-evolution and the hierarchy of phylogenetic classification across the bacterial tree-of-life (*13*). This result motivated investigating whether there was a relationship between the principal components of covariation amongst existing taxa and the pattern of diversifications by which the taxa were created. To do this, we generated *in silico* trajectories of diversification where we knew the precedent branching structure and examined the structure of covariation amongst the resulting taxa using a technique called Singular Value Decomposition (SVD). In this section we (i) describe SVD, (ii) describe how we related the output of SVD with paths of diversification that resulted in taxonomic diversity, and (iii) how we created a tree of taxonomic relatedness.

*1.1 A description of Singular Value Decomposition*

Singular Value Decomposition (SVD) is a matrix factorization method that is a generalization of Principal Components Analysis (PCA). In the main text, we use the term ‘principal components’; we note that principal components are also called of ‘spectral components’, ‘modes of variation’, or ‘eigenmodes’ in the literature. All of these terms describe the same mathematical concept as discussed below.

In general, SVD factorizes a real matrix **M** into three matrices according to the following equation:

$$M = U\Sigma V^T \quad (1)$$

In equation (1), **U** is termed the left-singular vector (LSV) matrix, **Σ** is a diagonal matrix of singular values, and **V** is termed the right-singular vector matrix. If **M** is a matrix of *n* systems (rows) described by *m* features (columns) where $n < m$; **U** is an $n \times n$ matrix where rows are systems, columns are LSVs, and each entry is the contribution of a given system to an LSV; **Σ** is an $n \times m$ matrix where the $k^{th}$ diagonal entry is the $k^{th}$ singular value and all off-diagonal entries are 0; and **V** is an $m \times m$ matrix where rows are features, columns are RSVs, and each entry is the contribution of a given feature to an RSV. $\mathbf{V}^\mathbf{T}$ in (1) is the transpose of **V**. A ‘spectral component’ is the axis specified by the $k^{th}$ singular value and is the same as a ‘principal component' from PCA, or 'eigenmode', or 'mode of variation'. The relationship between SVD and PCA is that PCA is performed only on either the rows or the columns of **M**. Therefore, matrices **U** and **Σ** can be multiplied together to form **P** which are exactly the principal components of matrix **M**.

$$P = U\Sigma \tag{2}$$

*1.2 Relating spectral properties of taxonomic covariation with diversification histories.*

The diversification trajectory shown in **Fig. S3A** is a representative of the toy models we used to conduct our analysis. In the example in **Fig. S3A**, each taxon in the alignment is defined by a 'genotype' comprised of fourteen features that are either a '1' or a '0'; each taxon is created from a series of three sequential diversification events. Collectively, the alignment of taxa represents extant diversity. The ancestral root is defined as a genotype of all '1'. The first layer of diversification from the ancestral root is defined by two separate mutations in positions 1 and 2. The second layer of diversification mutates positions 3 through 6 to create sub-populations. The third layer of diversification mutates positions 7 through 14 to create the extant diversity of taxa—eight taxa in total with diverse genotypes.

SVD on the alignment of taxa yielded eight spectral components (**Fig. S3B**). The extent to which each taxon contributes, or 'projects' onto each spectral component is shown in **Fig. S3C**. When visualizing the contribution of each taxon onto each spectral component, we observe a key finding. The taxa arising from a common broad layer of diversification contribute similarly to the first two spectral components while those arising from common finer layers of diversification (the second and third diversifications) contribute similarly to deeper spectral components. Thus, the statistical patterns contained in deeper spectral components are essential for defining fine-grained differences between taxa.

We translated our finding into a mathematical entity by computing the 'spectral distance' between two taxa. The spectral distance between two taxa, $\boldsymbol{i}$ and $\boldsymbol{j}$, on spectral component k is

$$SD_{ij}^{k} = \left|P_i^k - P_j^k\right| \tag{3}$$

where $\boldsymbol{P_i^k}$ is the projection of taxa $\boldsymbol{i}$ onto spectral component *k* and $\boldsymbol{P_j^k}$ is the contribution of taxa $\boldsymbol{j}$ onto spectral component *k*. We show an example of a pattern of spectral distances in **Fig. S3D** where taxon 'a' is the reference. Taxon 'a' and all other taxa share the same projection onto the first spectral component. As such, the spectral distance between 'a' and any other taxa is zero at spectral component 1. However, at spectral component 2, 'a' continues to share the same projection as taxa 'b', 'c', and 'd', but taxa 'e', 'f', 'g', and 'h' have a different projection onto spectral component 2. Therefore, the spectral distance between 'a' and 'e', 'f', 'g', and 'h' is non-zero at spectral component 2 but still zero for 'b', 'c', and 'd'.

We next defined the 'cumulative spectral distance' between two taxa from the first to $k^{th}$ spectral component as

$$SD_{ij}^{1:k} = \left|P_i^k - P_j^k\right| + \sum_{r=1}^{k-1}\left|P_i^r - P_j^r\right| \tag{4}$$

where *r* denotes the index of each spectral component 'shallower' than spectral component *k* (i.e., each spectral component with an associated singular value greater than the $k^{\text{th}}$ singular value). The cumulative spectral distance pattern of all pairs of taxa with taxa 'a' as one member of the pair is shown in **Fig. S3D**. Computing the cumulative spectral distance across all spectral components for all pairs of taxa illustrated a distinct tree-like pattern of partitioning between taxa.

A key finding from **Fig. S3D** was that groups of sequential spectral components collectively described different layers of precedent diversifications. For instance, in the example shown in **Fig. S3A**, spectral components 5 through 8 harbored the same amount of data-variance and collectively described the third layer of diversification from the ancestral root. Similarly, spectral components 3 through 4 harbored the same amount of data-variance and described the second layer of diversification from the ancestral root. Spectral components 1 and 2 independently described the ancestral root and the first layer of diversification respectively. This result motivated 'renormalizing' the spectral components into groups of spectral components harboring the same percent data-variance (**Fig. S3E**, left panel). Spectral component groups were defined based on the natural $\log_{10}$ difference between subsequent singular values. The rationale behind this choice was to group together spectral components that are relatively similar in their measure of percent-variance explained. To reduce effects at very small singular values, a pseudocount of 1 is added to each singular value. The difference between each subsequent singular value is expressed as

$$\Delta_k = \ln\left(\Sigma_{k-1,k-1} + 1\right) - \ln\left(\Sigma_{k,k} + 1\right) \tag{5}$$

Then, the $k^{th}$ component was chosen to start a new spectral group only if the difference between the $(k-1)^{th}$ and $k^{th}$ component was greater than a manually chosen threshold θ,

$$K = \{k \mid \Delta_k > \theta\} \tag{6}$$

For our *in-silico* toy models, the threshold $\theta$ was $0$ and therefore any drop in explained variance was defined as a new group of spectral components. For real biological data described in the main text, the threshold θ was chosen as 1.5 times the third quantile of these natural log differences ($\theta = 1.5 \times Q3(\Delta)$) as an approximation for selecting the only the largest drops in explained variance. The spectral distance computed across spectral component groups was defined as

$$SD_{ij}^{G} = \sum_{g \in G} |P_i^g - P_j^g|_{l2} \tag{7}$$

where *G* is the total set of spectral component groups, *g* is a specific spectral component group within *G*, and $|\cdot|_{l2}$ denotes the $l_2$ norm also known as the Euclidean distance.

*1.3 Creating a tree of relatedness from covariation structure of taxa*

Using **eq. (7)**, inferred trees of taxonomic relatedness ('Spectral Trees' in the main text) are generated using four steps.

*Step 1*: A reference pairwise spectral distance matrix **SD** is created from **eq. (7)** for all pairs of taxa comprising a matrix **M**.

*Step 2*: A reference tree $\mathbf{ST_{ref}}$ is generated via hierarchical clustering with average linkage, also known as the UPGMA (unweighted pair group method with arithmetic mean) method of phylogenetic tree building (*17*).

*Step 3*: A set of 100 bootstrap trees is generated using steps 1-2. For each bootstrap, we replace the original matrix **M** with a bootstrap matrix $\mathbf{M_{boot}}$ by sampling features (columns) with replacement to maintain the original dimensions of **M**. This procedure first generates a pairwise distance matrix $\mathbf{SD_{boot}}$ from the matrix $\mathbf{M_{boot}}$, and then generates a tree $\mathbf{ST_{boot}}$ using the UPGMA algorithm.

*Step 4*: The reference tree $\mathbf{ST_{ref}}$ and the bootstrap trees are then compared with transfer bootstrap expectation (TBE) as described in Lemoine *et al* (*18*). TBE ranges from 0 to 1 where 0 indicates no similar branches in any bootstrap tree, and 1 indicates the exact branch was found across all bootstrap trees.

The result of implementing these four steps generates a rooted Spectral Tree where each branch of the tree has an associated measure of support as defined by TBE. The Spectral Tree associated with the alignment in **Fig. S3A** is shown in **Fig. S3E** (right panel). We found that the Spectral Tree in **Fig. S3E** closely matched existing approaches of phylogenetic inference spanning maximum likelihood and Bayesian methods (**Fig. S3F**).

**2. A mathematical analysis of why spectral factorization reveals a hierarchy of relatedness.**

Given the multitude of possible factorization strategies, we sought to understand why spectral factorization in particular, i.e. SVD, reveals hierarchical scales of relatedness.

*2.1 The relationship between system similarity and the characteristic polynomial*

Spectral factorization uses the following equation to define spectral components of data-variance:

$$\det(D - \lambda I) = 0 \tag{8}$$

where 'det' is a mathematical function called the determinant, **D** is a matrix, **I** is the identity matrix of the same size as **D**, and $\lambda$ are solutions to **eq. (8)** known as 'eigenvalues'. Another form of **eq. (8)** is called the 'characteristic polynomial':

$$\det(D - \lambda I) = \lambda^n - c_{n-1}\lambda^{n-1} + c_{n-2}\lambda^{n-2} - c_{n-3}\lambda^{n-3} \ldots = 0 \tag{9}$$

Solving the roots of **eq. (9)** determines the eigenvalues of **D**. When performing spectral factorization on an ensemble of systems, **D** is a measure of distance (i.e. a covariance matrix) computed from a matrix of an ensemble of systems **M**. Relating how the structure of **D** is reflected in **eq. (9)** becomes increasingly complicated as the number of terms in **eq. (9)** increases (i.e. the number of systems in **M** increases). Thus, to interrogate how the relatedness of systems is manifest through **eq. (9)**, we created a simple ensemble comprised of three systems defined by a vector of ones and zeros–A, B, and C–where A and B shared complete similarity to each other and the similarity of C to A and B was tuned from sharing no similarity to complete similarity (**Fig. S4A**). The similarity matrix for each case is shown in **Fig. S4B** and provides a quantitative parameter to define the changing degree of relatedness. We term this variable $\gamma$; it is the off-diagonal terms of the first row and first column of the similarity matrices shown in **Fig. S4B**. We can write the determinant of the similarity matrices in general form:

$$\det\begin{pmatrix} \langle C|C\rangle - \lambda & \gamma & \gamma \\ \gamma & \langle B|B\rangle - \lambda & s \\ \gamma & s & \langle A|A\rangle - \lambda \end{pmatrix} = 0 \tag{10}$$

where *s* is the similarity between system A and B and is constant in our simple model of systems A, B, and C. Expanding **eq. (10)**, we get

$$(\langle C|C\rangle - \lambda)\begin{vmatrix} \langle B|B\rangle - \lambda & s \\ s & \langle A|A\rangle - \lambda \end{vmatrix} - \gamma\begin{vmatrix} \gamma & s \\ \gamma & \langle A|A\rangle - \lambda \end{vmatrix} - \gamma\begin{vmatrix} \gamma & \langle B|B\rangle - \lambda \\ \gamma & s \end{vmatrix} \tag{11}$$

From **eq. (11)**, we see that if $\gamma = 0$, all the terms that compare the similarity of C to A and B go to zero and the only remaining term reflects the similarity of C onto itself, separating the determinant of A and B. Moreover, from **eq. (11)**, we see that $\gamma$ is only in terms with a single $\lambda$. Thus, the only term in the characteristic polynomial dependent on $\gamma$ is the first order term of $\lambda$. Given the similarity matrices **Fig. S4B**, the characteristic polynomial including $\gamma$ for our model is

$$\lambda^3 - 9\lambda^2 + (18 - 2\gamma^2)\lambda = 0 \tag{12}$$

Substituting in the various values of γ defined in **Fig. S4B** gives different instances of the characteristic polynomial

- Unrelated ($\gamma = 0$): $\lambda^3 - 9\lambda^2 + 18\lambda$
- Related ($\gamma = 1$): $\lambda^3 - 9\lambda^2 + 16\lambda$
- Related ($\gamma = 2$): $\lambda^3 - 9\lambda^2 + 10\lambda$
- Identical ($\gamma = 3$): $\lambda^3 - 9\lambda^2$

Thus, as C becomes more related to A and B, the coefficient of the first order λ term (the third term in the characteristic polynomial) tends to zero.

<u>*2.2 Relating system similarity with pattern of eigenvalues*</u>

What is the consequence of the first order coefficient tending towards zero as C becomes more similar to A and B on the pattern of resulting eigenvalues? We find that as the similarity of C towards A and B increases, the two eigenvalues computed from the similarity matrices change in opposite directions: $\lambda_1$ becomes more positive while $\lambda_2$ becomes more negative (**Fig. S4C**). We find this relationship is a natural consequence of solving the characteristic polynomial. To show this, we start with **eq. (12)** and factor out a root and power of λ.

$$\boldsymbol{(\lambda - 0)\left(\lambda^2 - 9\lambda^1 + \left(18 - 2\gamma^2\right)\right) = 0} \tag{13}$$

In our case study, the factored-out root is the third eigenvalue and is always equal to zero because A and B are identical. Now, we use the quadratic formula to solve for the roots on the remaining 2nd order polynomial:

$$\frac{-b \pm \sqrt{b^2 - 4ac}}{2a} \rightarrow \frac{9 \pm \sqrt{81 - 4(18 - 2\gamma^2)}}{2} = \frac{9 \pm \sqrt{8\gamma^2 + 9}}{2} \tag{14}$$

**Eq. (13)** can be rewritten as

$$(\lambda - 0)\left(\lambda - \frac{9 - \sqrt{8\gamma^2 + 9}}{2}\right)\left(\lambda - \frac{9 + \sqrt{8\gamma^2 + 9}}{2}\right) = 0 \tag{15}$$

Thus, the two non-zero roots diverge from each other as a function of γ. When γ is 0 the roots are {0, +3, +6}; when γ is 3 the roots are {0, 0, +9}. Importantly, this result shows that by mathematical definition, the second eigenvalue will only equal zero when A, B, and C are identical to each other. Any degree of dissimilarity between C compared to A and B will manifest as a non-zero second eigenvalue, and therefore two non-zero spectral components. Critically, this relationship is entirely independent of the percent-variance harbored by the second spectral component. Thus, this case study shows how spectral components harboring a minority of data-variance beyond what is typically considered signal encode important information regarding system differences by mathematical definition.

<u>*2.3 Relating system similarity with information content contained in eigenvectors*</u>

We sought to understand the relationship between defining system relatedness and order of eigenvector in the eigenspectrum. We computed the contribution of systems A, B, and C onto eigenvectors $v_1$ and $v_2$, defined by $\lambda_1$ and $\lambda_2$ respectively. For eigenvector $v_1$, we found that the contribution of A and B is relatively constant while the contribution of C rapidly changes from zero and asymptotically reaches the same constant as A and B. In contrast, for eigenvector $v_2$ we found that the contribution of C is relatively constant while the contribution of A and B rapidly changes from zero to asymptotically reach a constant value away from that of C **(Fig. S4D)**. Thus, eigenvector $v_1$ defines the similarity between A, B, and C, whereas eigenvector $v_2$ defines the difference between A, B, and C. But these results motivated a key mathematical question: how does varying the similarity of C to A and B have different effects on eigenvector $v_1$ compared to eigenvector $v_2$?

The eigenvector equation for the similarity matrices shown in **Fig. S4B** is

$$\begin{bmatrix} \langle C|C\rangle - \lambda & \gamma & \gamma \\ \gamma & \langle B|B\rangle - \lambda & s \\ \gamma & s & \langle A|A\rangle - \lambda \end{bmatrix} \begin{bmatrix} x_C \\ x_B \\ x_A \end{bmatrix} = \begin{bmatrix} 0 \\ 0 \\ 0 \end{bmatrix} \tag{16}$$

We are interested in how the contributions of each system ($\{x_A, x_B, x_C\}$) change with respect to the similarity parameter γ. To address this question, we computed the partial derivative of the contribution of each system onto either eigenvector $v_1$ or $v_2$ with respect to γ. Below, we describe this calculation in detail for each system.

System C:

From **eq. (16)**, we find that the way C is related to γ is through the following equation

$$(\langle C|C\rangle - \lambda)x_C + \gamma x_B + \gamma x_A = 0 \tag{17}$$

Isolating $x_C$ gives

$$(\langle C|C\rangle - \lambda)x_C + \gamma x_B + \gamma x_A = 0 \tag{18}$$

$$(\langle C|C\rangle - \lambda)x_C = -(\gamma x_B + \gamma x_A) \tag{19}$$

$$x_C = \frac{-\gamma(x_B + x_A)}{(\langle C|C\rangle - \lambda)} \tag{20}$$

So

$$x_C = \frac{\gamma(x_B + x_A)}{(\lambda - \langle C|C\rangle)} \tag{21}$$

What are $x_B$ and $x_A$? Assuming an expansion around the case where C is unrelated to A and B, **eq. (16)** becomes

$$\begin{bmatrix} 3-\lambda & 0 & 0 \\ 0 & 3-\lambda & 3 \\ 0 & 3 & 3-\lambda \end{bmatrix} \begin{bmatrix} x_C \\ x_B \\ x_A \end{bmatrix} = \begin{bmatrix} 0 \\ 0 \\ 0 \end{bmatrix} \tag{22}$$

The first root is λ = 6 when γ is zero. This makes **eq. (22)**

$$\begin{bmatrix} -3 & 0 & 0 \\ 0 & -3 & 3 \\ 0 & 3 & -3 \end{bmatrix} \begin{bmatrix} x_C \\ x_B \\ x_A \end{bmatrix} = \begin{bmatrix} 0 \\ 0 \\ 0 \end{bmatrix} \tag{23}$$

To solve for $x_C$,

$$-3x_C + 0x_B + 0x_A = 0 \tag{24}$$

Thus, $x_C$ is equal to zero. The equations for $x_B$ and $x_A$ work together to show that they can equal any real number so long as $x_A = x_B$.

$$3x_A - 3x_B = x_A - x_B = 0 \tag{25}$$

To make $[x_C, x_B, x_A]^t$ a unitary vector (i.e., with length equal to 1), we set $x_A = x_B = \frac{1}{\sqrt{2}}$.
The second root is λ = 3. So then, **eq. (16)** becomes

$$\begin{bmatrix} 0 & 0 & 0 \\ 0 & 0 & 3 \\ 0 & 3 & 0 \end{bmatrix} \begin{bmatrix} x_C \\ x_B \\ x_A \end{bmatrix} = \begin{bmatrix} 0 \\ 0 \\ 0 \end{bmatrix} \tag{26}$$

To solve for $x_C$,

$$0x_C + 0x_B + 0x_A = 0 \tag{27}$$

Thus, $x_C$ can be any value. The equations for $x_B$ and $x_A$ show that they must both equal zero.

$$0x_C + 0x_B + 3x_A = x_A = 0 \tag{28}$$

$$0x_C + 3x_B + 0x_A = x_B = 0 \tag{29}$$

To make $[x_C, x_B, x_A]^t$ a unitary vector (i.e., with length equal to 1), we set $x_A = x_B = 0$ and $x_C = 1$. So, overall $x_B$ and $x_A$ are $\frac{1}{\sqrt{2}}$. for $v_1$ and 0 for $v_2$. Plugging these values of $x_B$ and $x_A$ into **eq. (21)** for $v_1$ gives

$$x_C = \frac{\gamma(x_B + x_A)}{\lambda_1 - \langle C|C\rangle} = \frac{\gamma\left(\frac{1}{\sqrt{2}} + \frac{1}{\sqrt{2}}\right)}{\lambda_1 - \langle C|C\rangle} = \frac{2\gamma}{\sqrt{2}(\lambda_1 - \langle C|C\rangle)} \tag{30}$$

$$x_C = \frac{\sqrt{2}\gamma}{\lambda_1 - \langle C|C\rangle} \tag{31}$$

For $v_2$

$$x_C = \frac{\gamma(x_B + x_A)}{\lambda_2 - \langle C|C\rangle} = \frac{\gamma(0 + 0)}{\lambda_2 - \langle C|C\rangle} \tag{32}$$

$$x_C = 0 \tag{33}$$

Thus, for $v_1$

$$\frac{\partial x_C}{\partial \gamma} = \frac{\sqrt{2}\gamma}{\lambda_1 - \langle C|C\rangle} \tag{34}$$

and for $v_2$

$$\frac{\partial x_C}{\partial \gamma} = 0 \tag{35}$$

Following the same procedure described from **eqs. (17)** to **(35)** for $x_C$ on $x_B$ and $x_A$, we find that for $v_1$

$$\frac{\partial x_B}{\partial \gamma} = \frac{\partial x_A}{\partial \gamma} = 0 \tag{36}$$

Whereas for $v_2$

$$\frac{\partial x_B}{\partial \gamma} = \frac{1}{\lambda_2 - \langle B|B\rangle} \tag{37}$$

$$\frac{\partial x_A}{\partial \gamma} = \frac{1}{\lambda_2 - \langle A|A\rangle} \tag{38}$$

In sum, the way that the contributions of each system onto eigenvectors 1 and 2 change with respect to $\gamma$ is summarized in the table below:

| | $\frac{\partial x_C}{\partial \gamma}$ | $\frac{\partial x_B}{\partial \gamma}$ | $\frac{\partial x_A}{\partial \gamma}$ |
|---|---|---|---|
| ***Eigenvector*** *$v_1$* | $\frac{\sqrt{2}}{\lambda_1 - \langle C\|C\rangle}$ | 0 | 0 |
| ***Eigenvector*** *$v_2$* | 0 | $\frac{1}{\lambda_2 - \langle B\|B\rangle}$ | $\frac{1}{\lambda_2 - \langle A\|A\rangle}$ |

Thus, we find that in the limit of a small increase in similarity of C to A and B, the change in the contribution of C to eigenvector $v_1$ is a constant while the that of A and B is zero. In contrast, the change in the contribution of C to eigenvector $v_2$ is zero while that of A and B is a constant. Because $\lambda_1$ and $\lambda_2$ trend in opposite directions per **eq. 15**, the contribution of C to eigenvector $v_1$ smoothly tends towards that of A and B thereby defining the extent of similarity between systems while the contributions of A and B to eigenvector $v_2$ smoothly tend away from that of C thereby defining the extent of dissimilarity between systems. In totality, these results demonstrate that the mathematical structure of spectral factorization is inherently able to define scales of system relatedness and orthogonalizes this information into separate eigenvectors.

That is, by definition the shallowest eigenvectors define patterns of global similarity between systems while deeper eigenvectors renormalize system relationships to define nested differences amongst global similarities.

*2.3 Exploring the accuracy of Spectral Trees in capturing hierarchies of taxonomic relatedness across a broad set of parameters*

The mathematical analysis described in Sections 2.1 to 2.3 suggested that defining accurate hierarchies of relatedness between taxa from using spectral factorization is robust to (i) size of the alignment and (ii) number of features describing each system in the alignment. To test this idea, GoTree v 0.4 (https://github.com/evolbioinfo/gotree) was used to create reference, 'ground-truth', trees of taxa (*19*). We used GoTree to create 7 separate trees comprising either 16, 32, 64, 128, 256, 512, or 1,024 leaves. Each tree was then input into SeqGen v 1.3 (https://github.com/rambaut/Seq-Gen) which produced multiple sequence alignments (MSAs) where rows were leaves and columns were features describing the leaves (*20*). SeqGen uses a Markov process considering the branching pattern of the tree to create a vector of features for each leaf. Elements of the vector are the characters 'A' and 'T', and the Markov process uses uniform probabilities to flip between these characters at each branch point in the tree. For each of the 7 trees, we generated 7 MSAs with SeqGen, where each MSA contained either 16, 32, 64, 128, 256, 512, and 1024 features. Thus, our analysis spanned 49 total MSAs.

Spectral Trees were computed for each MSA. We then compared the topology of the resulting Spectral Trees against ground-truth defined by GoTree using an F-score—the harmonic mean of precision and recall. Precision between two trees is defined as the proportion of predicted branches in the Spectral Tree that are also in the 'true' tree. Recall is defined as the proportion of branches in the 'true' tree that are also in the Spectral Tree. F-score ranges between 0 to 1, where 1 indicates complete identity between the two trees and 0 indicates no commonality between the two trees.

Our results are shown in **Fig. S5**. We found that for the majority of the parameter space, the F-statistic was near 1. In the limit that the number of features was less than the number of taxa, the F-statistic was uniformly near 0. This distinction in F-statistic based on the parameter space arises from the scenario where the number of features is the limiting descriptor relative to the number of taxa in the alignment. The physical interpretation of this regime is that the number of features describing each system is substantially limited compared to the diversity of systems available for sampling. In this case, the information content of the set of features is 'overwritten' by the diversity of taxa, thereby erasing patterns of covariation originating from phylogenetic histories. A biological process that is consistent with this regime is if the recombination rate is extremely high relative to speciation events—a scenario that has been put forth as a plausible scenario for bacterial phylogenomic trends (*21–23*).

**3. Spectral Trees resolve trajectories of diversification with convergence.**

Our approach for creating Spectral Trees results in qualitatively distinct descriptions of relatedness relative to other methods of phylogenetic inference in cases of convergent processes. Convergent evolution involves two or more taxa possessing the same set of genomic traits through independent ancestral histories. These convergent histories vastly complicate phylogenetic inference because genomic diversity no longer increases in a predictable manner over evolutionary time. This problem is particularly evident when using single features (i.e. 'gene markers') to model ancestral distance because such an approach does not consider epistasis between genes and the contextual dependence of gene presence or absence on other genes. Methods using maximum likelihood and Bayesian phylogenetic inference using a Markov process typically assume site and mutation independence for practical reasons: accounting for every pairwise dependence between sites for large alignments defined by lots of features becomes computationally intractable (*8*, *14*–*16*, *24*).

A paradigmatic example of a trajectory with convergent processes is shown in **Fig. S6A**. The ancestral root ('F0 generation') is defined by a 'genotype' of eight 1's; there are three sequential sets of diversifications leading to 18 diverse taxa. After following the procedure outlined in **Section 1.3** of the Supplementary Text, we created a Spectral Tree of taxa (**Fig. S6B**,**C**). We found that the Spectral Tree correctly captured the generative set of diversifications spanning the F1 and F2 generations while application of other methods (FastMe and PhyML) did not (**Fig. S6C**). To better understand this result, we analyzed how features in the alignment were contributing to each spectral component. As an example, we considered position 5 in the alignment higlighted in **Fig. S6D**. Position 5 is a '0' for all taxa arising from the top branch of the 'F1' diversification event and is a '0' for one-third of taxa arising from the bottom branch of the 'F1' diversification event. Thus, two separate contexts, evolved independently, resulted in a '0' at position 5—an example of convergence. Using position 5 as a 'marker position' would therefore incorrectly group taxa arising from different histories together—a commonly encountered problem when considering each genomic feature in isolation of its genetic context (*25*).

A unique quality of our approach is that the Spectral Tree is constructed by considering all spectral components of variance, including those typically discarded as statistical noise. We sought to elucidate where information regarding the different generations of diversifications lay across the set of spectral components. We conducted this analysis by first defining sequential windows of spectral components across all nine spectral components (components 1 to 3, 2 to 4, …, 7 to 9). For each spectral window, we isolated the corresponding LSVs from the $U$ matrix defined by **eq. (1)**. This results in several sub-matrices defined by taxa on the rows, LSVs on the columns, and each entry being the contribution of each taxon onto each LSV. For each sub-matrix constructed from $U$, we computed the Pearson correlation between all pairs of taxa across the set of LSVs defined in the sub-matrix ('spectral correlations'). As a concrete example, for the first spectral window comprising spectral components 1 to 3, the $U$ submatrix is defined as taxa (rows) and the first three columns (LSV1 to LSV3) of the $U$ matrix. Then, to compute the spectral correlations between all pairs of taxa within LSVs 1 to 3, we computed the Pearson correlations between all pairs of rows in the sub-matrix. The result is a taxon-by-taxon spectral correlation matrix where each entry is the Pearson correlation measured between two taxa across spectral components 1 to 3. Defining all taxon-by-taxon spectral correlation matrices across all spectral windows creates a three-dimensional tensor, **R**, defined by taxa (rows), taxa (columns), and spectral windows (z-axis) where each entry in the tensor is the spectral correlation between two taxa within a spectral window. Separately, we created a second tensor, **G**, where rows are defined as taxa, columns are defined as taxa, the z-axis is each generation ('F0', 'F1', 'F2', or 'F3'), and entries in the tensor are a '1' if two taxa are grouped within the same cluster at a given generation or '0' if two taxa are not grouped within the same cluster at a given generation. We then computed the mutual information (MI) between each face of the R tensor with each of the G tensor. This computation interrogated the information shared between spectral correlations between taxa and shared generational history. The MI was calculated as

$$MI(r|G) = H(r) - \left(\frac{n_0}{N} H(r_0) + \frac{n_1}{N} H(r_1)\right) \tag{39}$$

where *H(r)* is a measure of entropy and is defined as

$$H(x) = \log_2(\Delta_{bw}) - \sum_b p(x_b) \log_2(p(x_b)) \tag{40}$$

$p(x_b)$ is the proportion of pairs that fall into a particular bin within a distribution of $x_b$ values; we use a bin-width of 0.01 to construct 200 bins across the distribution of correlation values ranging from -1 to 1; $r_1$ is the distribution of spectral correlations across taxonomic pairs that are descendants of the same ancestor; $n_1$ is the number of pairs in $r_1$; $r_0$ is the distribution of spectral correlations across taxonomic pairs that are not descendants of the same ancestor; $n_0$ is the count of those pairs within each bin; *N* is the total number of pairs. The meaning of this calculation is a measure of the extent to which knowing the distribution of spectral correlations within a spectral window between two taxa indicates the shared ancestral history of two taxa.

Our results showed that the shallowest set of principal components, spanning components 1 to 5, was enriched for information regarding clustering at the F1 Generation, but the deepest set of principal components, spanning components 4 to 8, was enriched for information regarding clustering at the F2 Generation (**Fig. S6E**). Using this information, we could then leverage a central property of SVD—its linearity—to isolate the statistical information in these different principal components. We can rewrite **eq. (1)** as

$$M = \sum_k \sigma_k u_k v_k^t \tag{41}$$

where $\sigma_k$ is the $k^{th}$ singular value and $u_k$ and $v^t_k$ are the $k^{th}$ left and right singular vectors. Each product that is being summed in **eq. (41)** is a rank 1 matrix because it produces a matrix that are scalar multiples of $v^t_k$. Using **eq. (41)**, we recreated the original alignment shown in **Fig. S6A** but only considering the information contained in principal components 5 to 8. This process is shown in **Fig. S6F**. Focusing on position 5 again, we found that the value of position 5 was adjusted in the recreated alignment reflecting the separate, nested contexts of diversification (**Fig. S6G**). Thus, by considering information contained across all principal components, the Spectral Tree accurately resolved both broad and context-dependent, finer patterns of diversification.

**4. A summary step-by-step workflow for computing a Spectral Tree for an ensemble of evolved systems.**

Assuming *N* systems are defined by *F* features, a matrix **M** is created where rows are systems and columns are features. The steps to create a Spectral Tree are:

(1) **M** is factorized using SVD [see **eq. (1)**].

(2) A cumulative spectral distance matrix across groups of spectral components (**SD**) is created from the output of SVD. [see **eq. (7)** and *Section 1.3*, Step 1].

(3) A bootstrapped tree with support values is created from hierarchical clustering of the **SD** matrix [see *Section 1.3*, Steps 2 through 4].

This rubric is applicable to any matrix of systems. In our work, the *N* systems comprise bacteria described by their gene content; the features of gene content collectively define *F*.

### Supplementary Figures

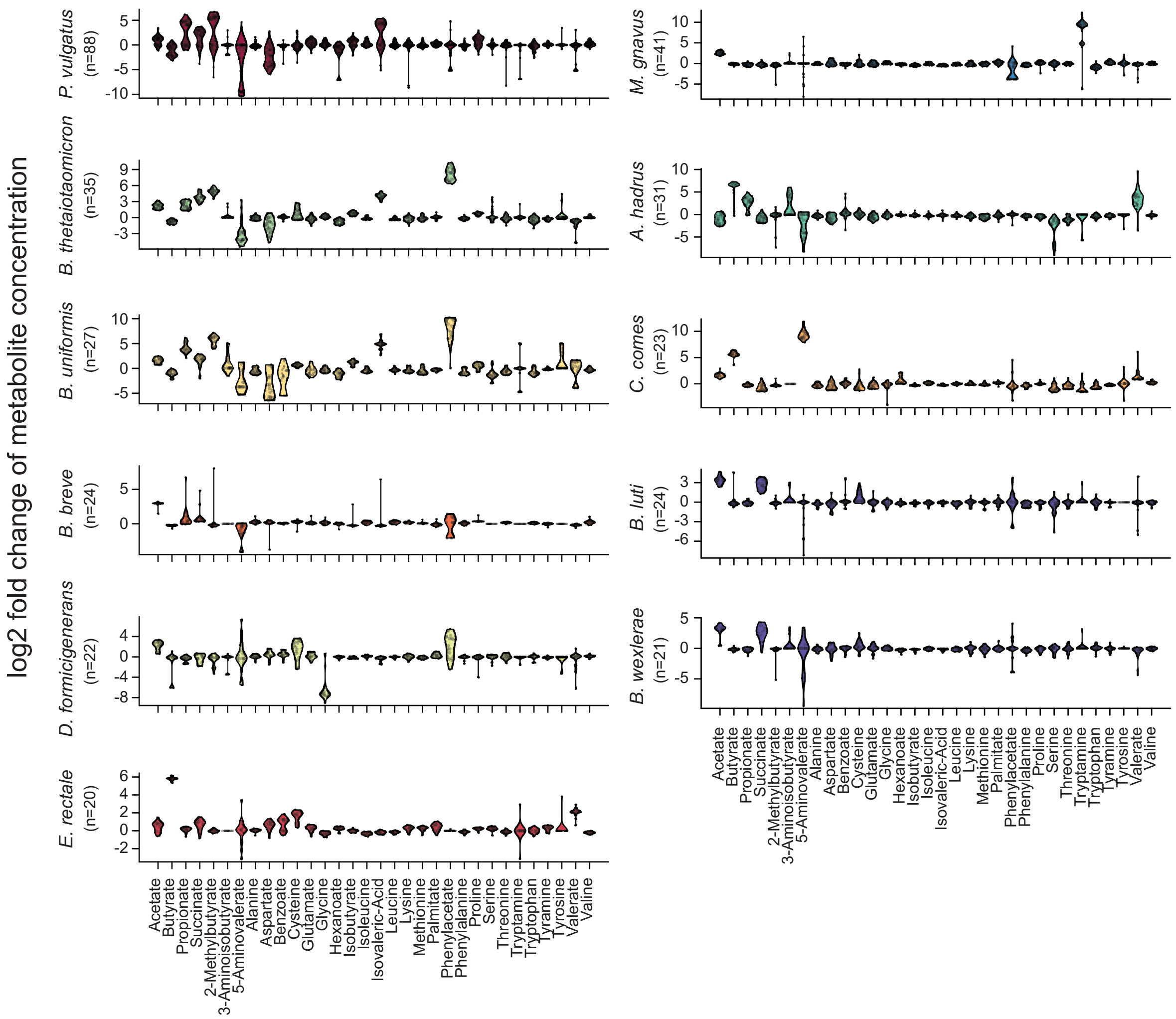


**Fig. S1.**
Metabolic variability across commensal strain bank. Strain-level metabolic variability across all species for which there were at least 10 strains. Each panel shows the log2 fold-change (y-axis) of each metabolic feature (x-axis) for a given species; number of strains within a species shown in parenthesis on the y-axis.

A.

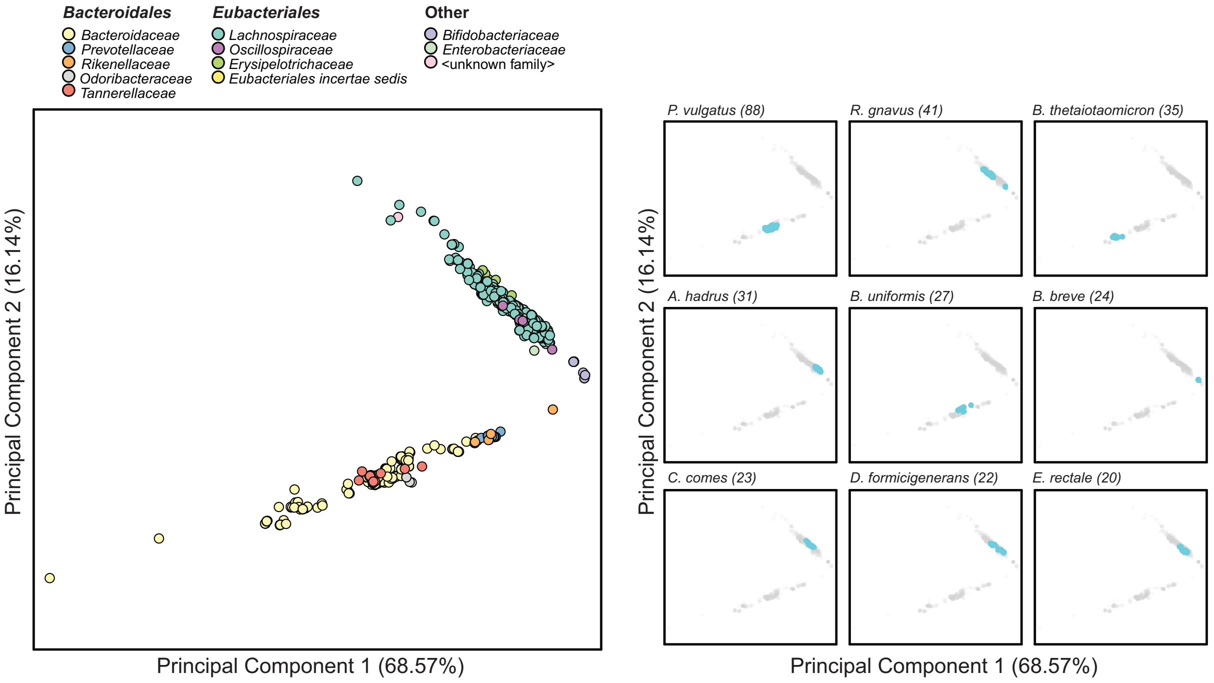


B.

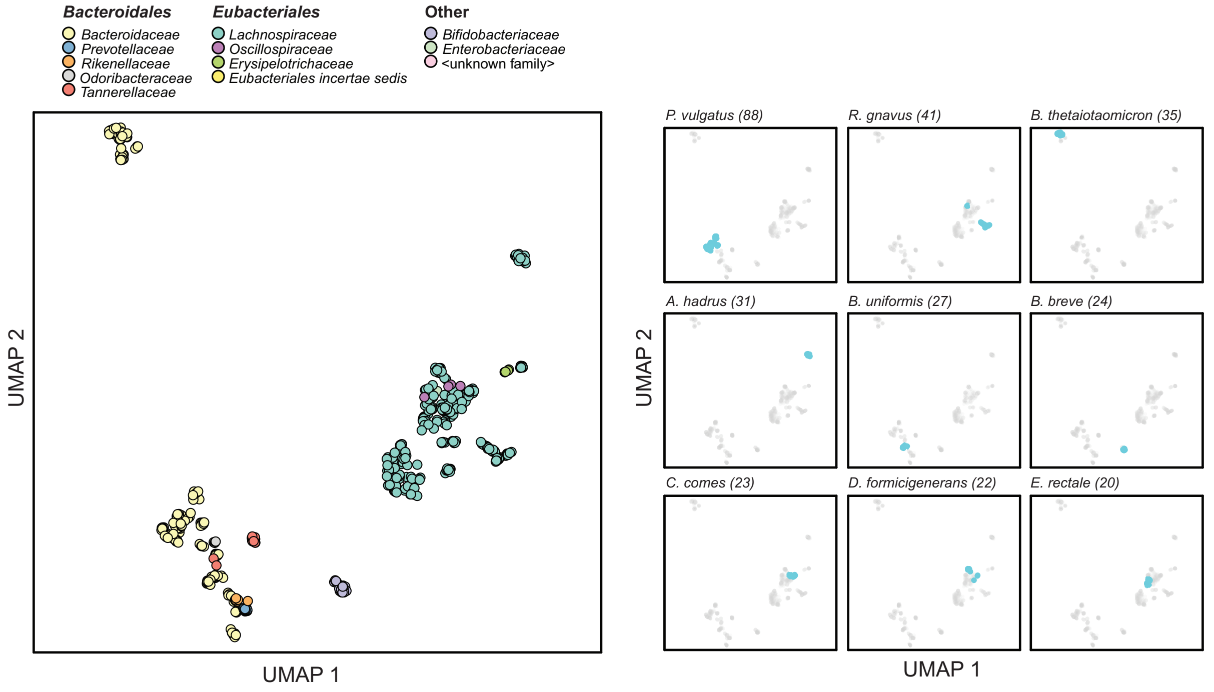


**Fig. S2.**
PCA and UMAP plot of commensal strain bank. Left panels; PCA (panel A) and UMAP (panel B) plots of commensal strain bank colored by ‘Family’ phylogenetic designation. Right panels; plots colored by specific strains. Number of strains designated in parenthesis.

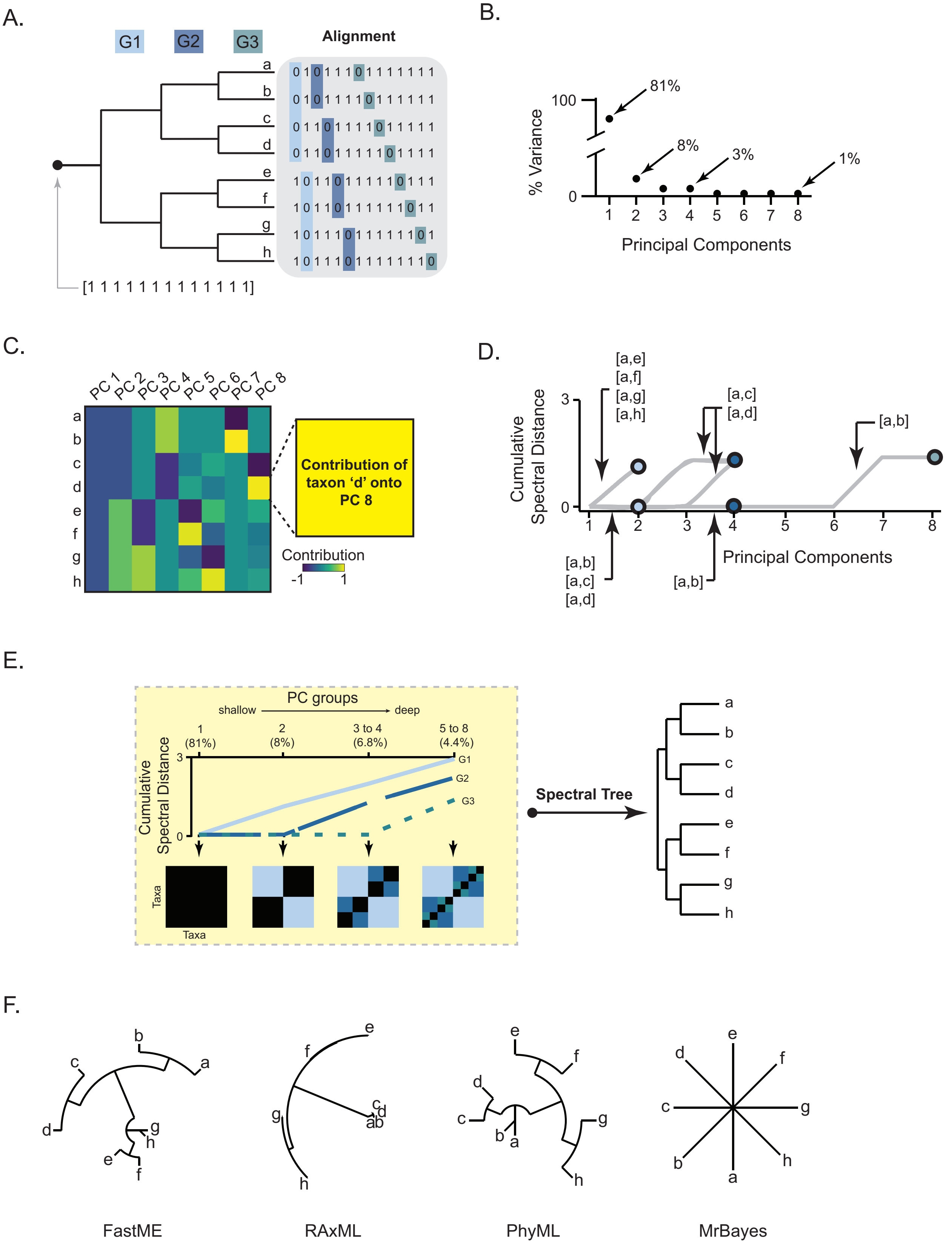

A.
G1
G2
G3
Alignment
[1 1 1 1 1 1 1 1 1 1 1 1 1]
B.
% Variance
Principal Components
81%
8%
3%
1%
C.
Contribution of taxon 'd' onto PC 8
Contribution
D.
Cumulative Spectral Distance
Principal Components
E.
PC groups
shallow
deep
Spectral Tree
Taxa
F.
FastME
RAxML
PhyML
MrBayes

**Fig. S3.**

Genomic distance inferred from covariation patterns. **(A)** An *in silico* model of sequential diversification. A ancestral root is defined by 13-bit string of ‘1’. Diversification through three generations creates an alignment of eight ‘taxa’. Colored bits in the alignment match the color of generation ‘G1’, ‘G2’, or ‘G3’ at which variation from a ‘1’ to ‘0’ was created. **(B)** Principal Components Analysis (PCA) of the alignment yields 8 principal components (PCs). Percent-variance of PC is shown. **(C)** Contributions of each taxon in the alignment in panel A to each PC. **(D)** Cumulative spectral distance (y-axis) for all pairs of taxa that include taxon ‘a’. The pattern of cumulative spectral distances resembles a tree-like distribution. **(E)** PCs are grouped together based on their percent variance. For each ‘PC group’, spectral distances are computed between all pairs of taxa. Taxa-taxa spectral distance matrices for each PC group are displayed; black to blue pixel colors indicate low to high spectral distances. This information is used to create a ‘Spectral Tree’. **(F)** Trees generated by various phylogenetic inference methods employed on the alignment in panel A.

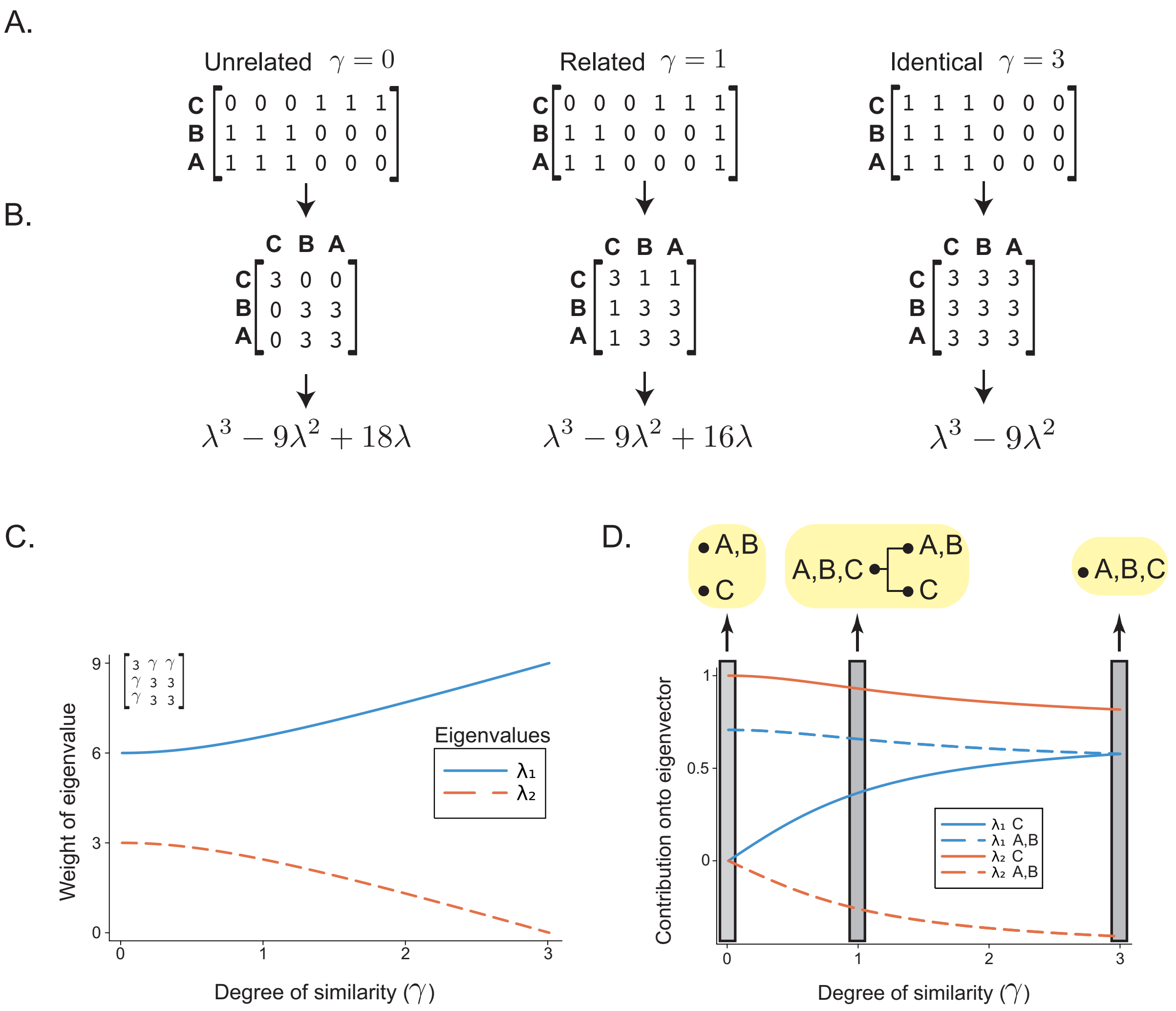


**Fig. S4.**
Spectral factorization defines hierarchies of relatedness between systems. **(A)** Three systems—A, B, and C—defined by six positions that can be either '0' or '1'. The similarity between system C and systems A and B ($\gamma$ value) is tuned. Examples of system C being unrelated (left), related (middle), and identical (right) to systems A and B are shown. **(B)** Similarity matrices between systems A, B, and C are shown with their associated characteristic polynomials. **(C)** Weight of eigenvalues $\lambda_1$ and $\lambda_2$ (y-axis) plotted against $\gamma$ (x-axis). **(D)** Contribution of each system onto eigenvectors 1 and 2 (y-axis, see color key) plotted against $\gamma$ (x-axis). Gray boxes delineate relatedness relationships between A, B, and C inferred from eigenvectors 1 and 2 at a given $\gamma$. The left box shows that C is completely dissimilar to A and B indicated by divergent eigenvectors 1 and 2. The middle box shows that A and B share a degree of similarity and dissimilarity with C indicated by a trending convergence in eigenvector 1 and trending divergence in eigenvector 2. The right-most box shows that A, B, and C are completely similar to each other indicated by a more convergent eigenvector 1 and more divergent eigenvector 2.

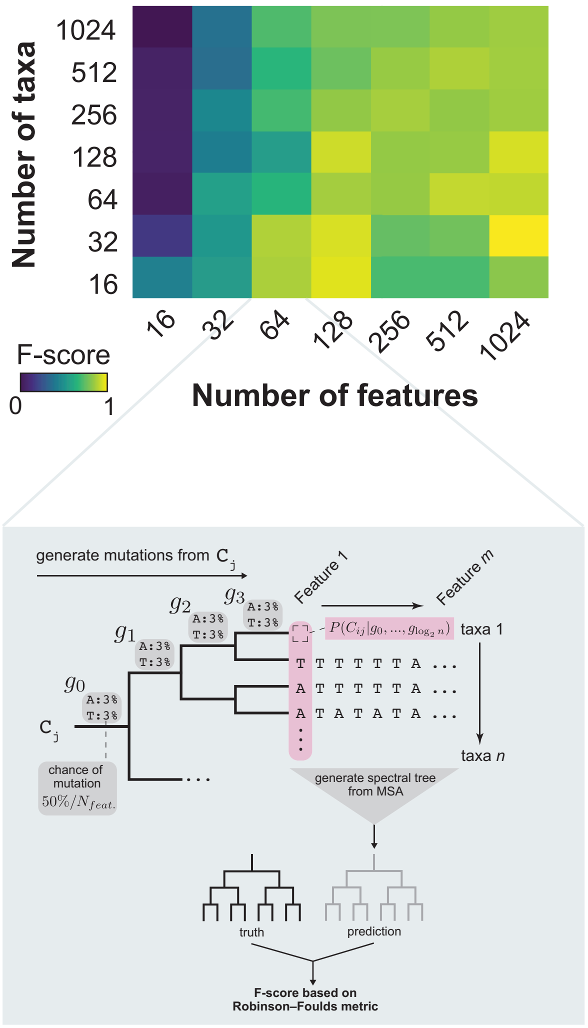


**Fig. S5.**
Accuracy of Spectral Tree architecture compared to ground truth patterns of diversification across many *in silico* models. Models varied by number of taxa (y-axis) and number of features defining each taxon (x-axis). F-scores (harmonic mean of precision and recall) were computed between the trajectory of sequential diversifications that created the diversity of resulting taxa and the architecture of the Spectral Tree. Inset shows an example of a single model. Within example model, an alignment of taxa is built from a tree and the Spectral Tree is compared to the ground truth of historical diversifications resulting in the alignment of taxa by an F-statistic (Robinson-Foulds metric).

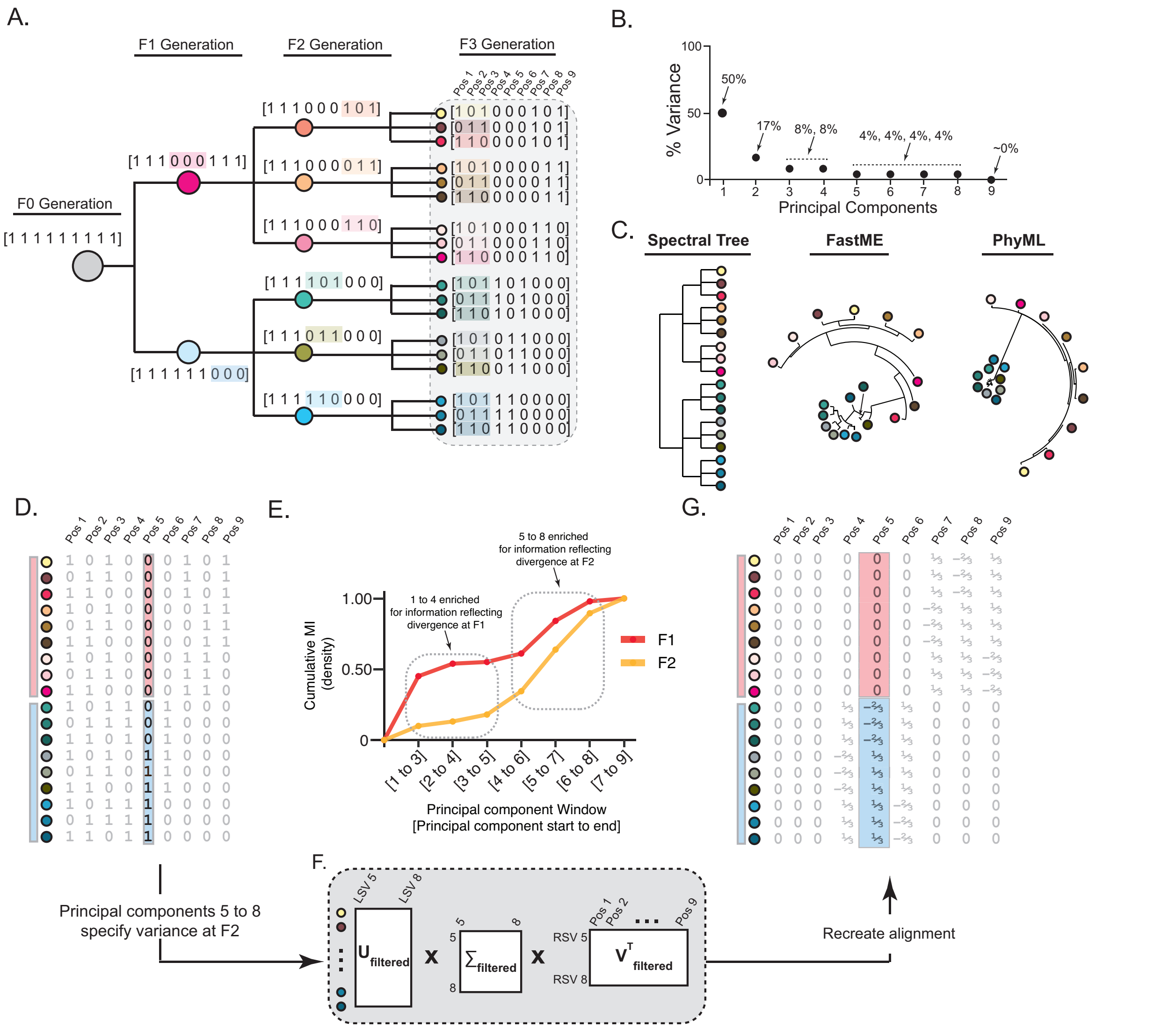


**Fig. S6.**
The Spectral Tree resolves patterns of convergent diversification trajectories. **(A)** Ancestral root (‘F0 Generation’) is defined by a 9-feature ‘genotype’ of ‘1’ and subject to three sequential diversification events (‘F1’, ‘F2’, and ‘F3’ generations) resulting in 18 taxa; features can be ‘1’ or ‘0’. Colored features in each generation correspond to variation from the previous generation. **(B)** Scree plot of 9 principal components (PCs) derived from SVD on alignment of taxa in F3 Generation from panel A. **(C)** Spectral Tree, tree resulting from FastME, and tree resulting from PhyML. **(D)** Alignment of taxa with position 5 highlighted; taxa are labeled by the group descend from in the F1 generation (red or blue bar). **(E)** Shared information (Cumulative Mutual Information (MI), y-axis) between clustering of taxa across PCs (x-axis) and whether taxa arise from F1 or F2 generations (color key). **(F,G)** Isolation of left singular vectors (LSV) 5 to 8 ($U_{filtered}$), singular values ($\Sigma_{filtered}$) 5 to 8, and right singular vectors (RSV) ($V^{T}_{filtered}$) 5 to 8 (panel F). Multiplication of filtered matrices in (F) produces a new 18 x 9 alignment containing only the information within PCs 5 to 8. Recreated alignment with position 5 highlighted; taxa are labeled by group they belong to at F1 generation (panel G).

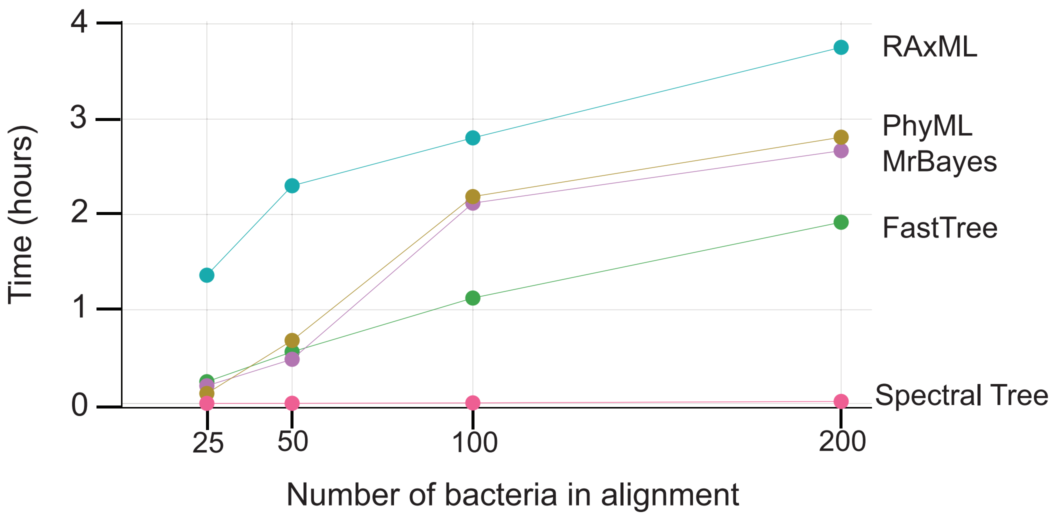


**Fig. S7.**
Comparison of compute times (y-axis) to create bootstrapped trees of bacterial relatedness for various methods of phylogenetic reconstruction. Times were computed in parallel on the University of Chicago Midway2 cluster, using a 2.4 GHz 7 core Intel E5-2680 v4 CPU with 16 GB of RAM per tree building task.

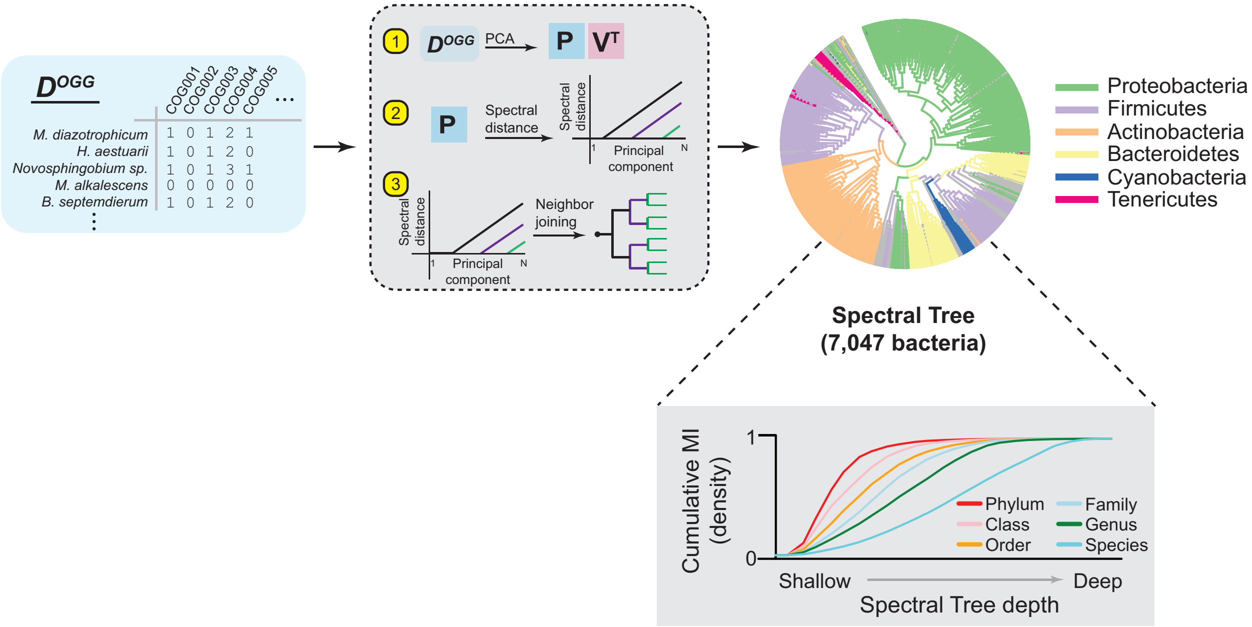


**Fig. S8.**
Workflow for computing the Spectral Tree of 7,047 non-redundant bacterial strains in the UniProt database. $\boldsymbol{D}^{OGG}$ is the matrix of 7,047 UniProt reference bacterial strains (rows) annotated by their OGG content (columns); each entry is the number of a specific OGG within a specific bacterial proteome. This matrix is then factorized using PCA and subject to the steps outlined in *Supplemental Text 1.3: Creating a tree of relatedness from covariation structure of taxa* to create a Spectral Tree. Inset shows the information shared between clusters of the Spectral Tree from shallow to deep (x-axis) and phylogenetic information (see color key). When a phylogenetic curve reaches a value of ‘1’, deeper clusters no longer group strains together by the labeled phylogenetic designation.

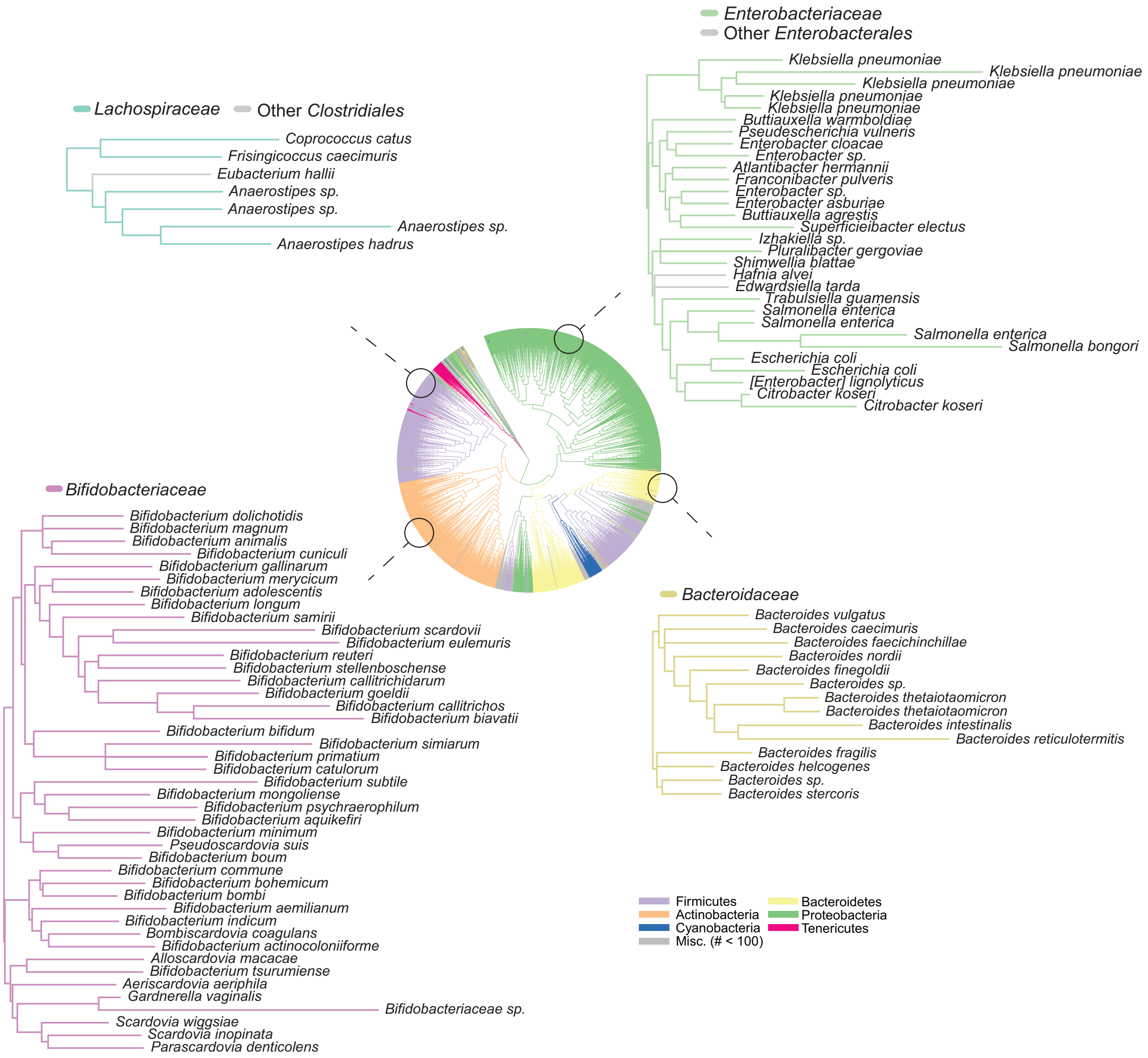


**Fig. S9.**
Spectral Tree of UniProt bacterial reference proteomes. Zoom-ins of Spectral Tree at specific bacterial Families are shown.

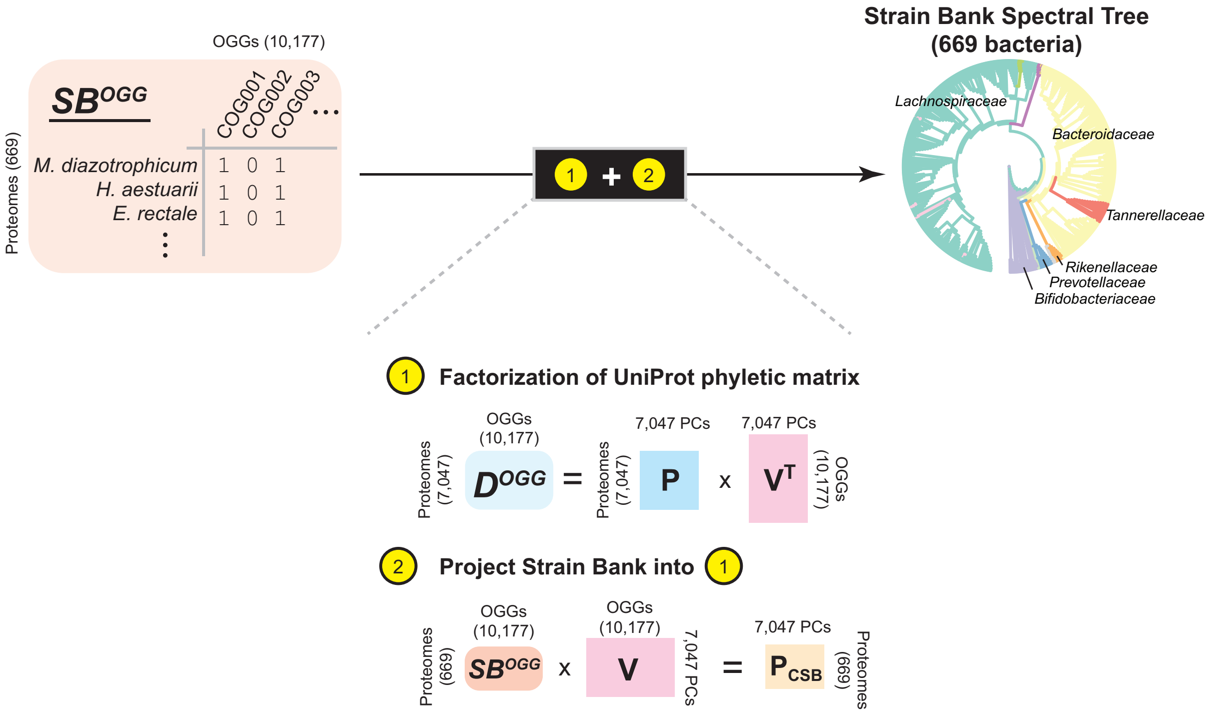


**Fig. S10.**
Workflow for projecting strains from the commensal strain bank into Spectral Tree. Whole-genome sequenced commensal strain bank strains are annotated by their Orthologous Gene Groups (OGGs) and aligned into a matrix (orange, $\boldsymbol{SB}^{OGG}$). These strains are then projected into the Spectral Tree by two steps. First, the phyletic matrix ($\boldsymbol{D}^{OGG}$) of UniProt reference proteomes (7,047 proteomes as rows and 10,177 OGGs as columns) is factorized into two matrices: **P** and $\mathbf{V}^T$. **P** defines the 7,047 principal components of bacterial co-evolution and $\mathbf{V}^T$ is the transpose of the matrix that defines the contribution of each OGG to each principal component. Second, $\boldsymbol{SB}^{OGG}$ is multiplied by **V** to compute the projection of each new strain onto the principal components defined by bacterial co-evolution across the UniProt reference proteomes. The result of implementing these two steps enables defining a tree of relatedness amongst the commensal strain bank in the context of co-evolutionary patterns reflected in the UniProt database.

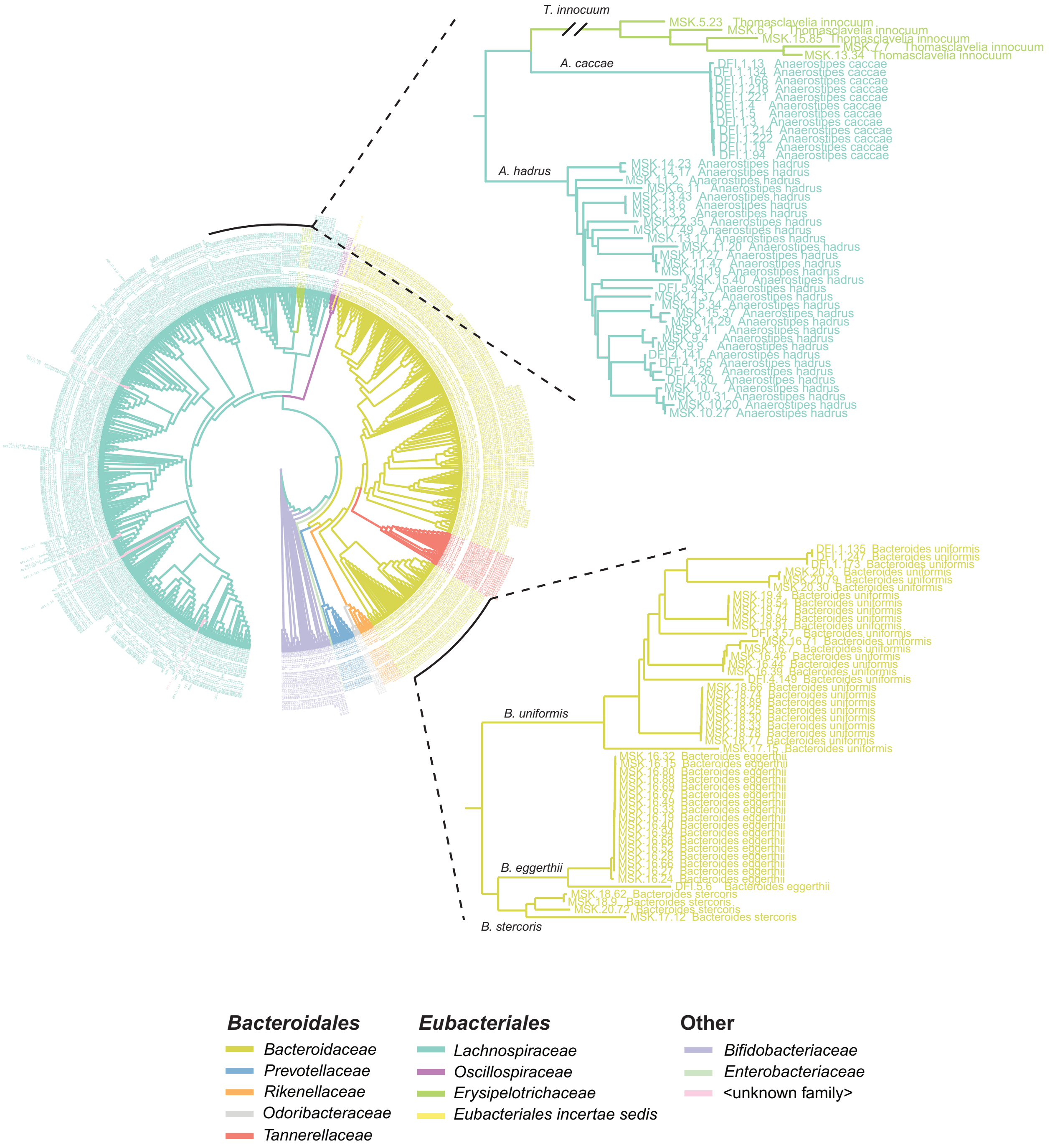


**Fig. S11.**
Tree of commensal strain bank strains defined by Spectral Tree. Strains are colored by ‘Family’ designation per NCBI taxonomy (see color key). Strain identities are shown, prefix indicates source of strain (MSK: Memorial Sloan Kettering Hospital; DFI: Duchossois Family Institute at University of Chicago).

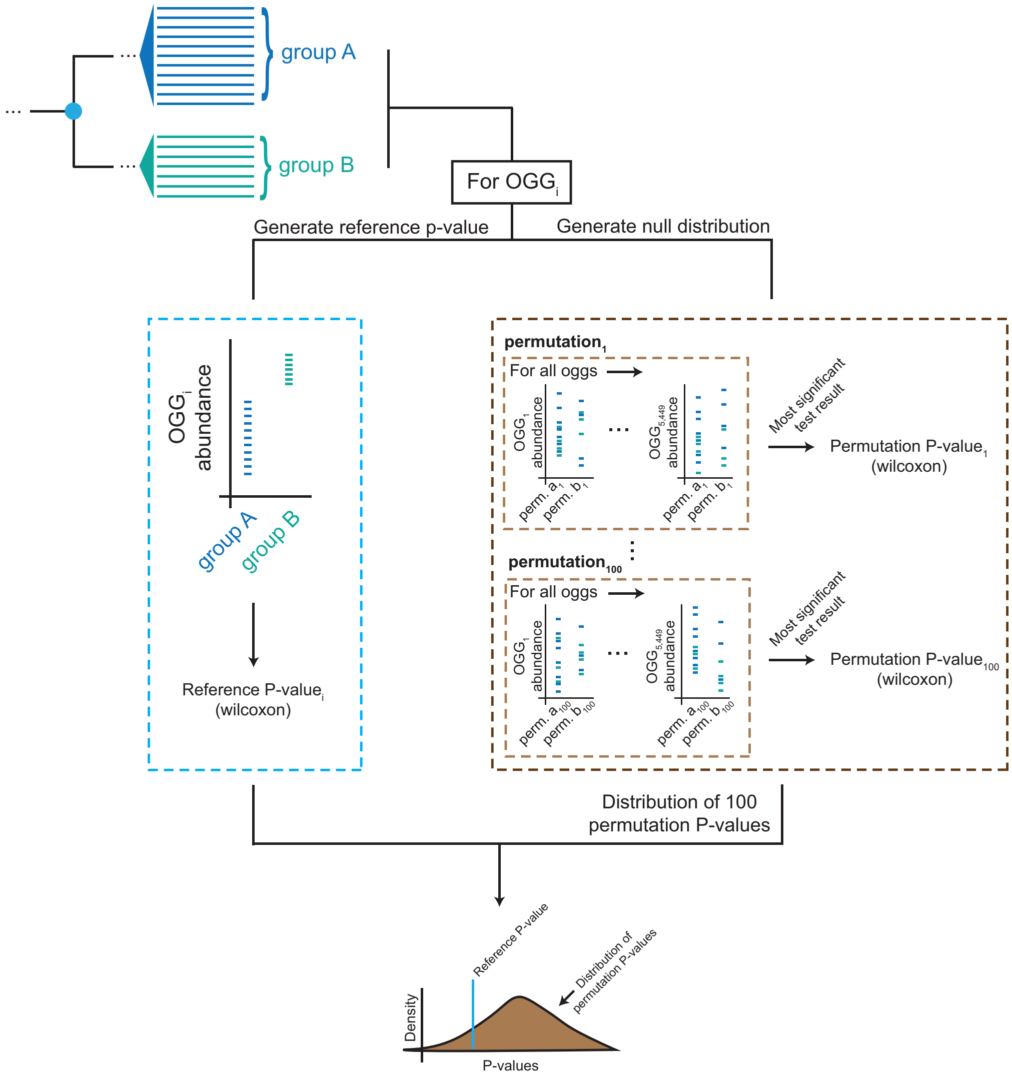


$$Q = \frac{N_{P_{perm}} < P_{ref}}{N_{P_{perm}}}$$

**Fig. S12.**
Workflow for determining the statistical significance of a differentially abundant OGG within the Spectral Tree. To determine whether a given OGG (OGG$_i$) is differentially abundant in a statistically significant way across two groups (‘group A’ and ‘group B’) that arise from a common node in the Spectral Tree, we first generate a ‘reference p-value’ and a ‘null distribution’. The reference p-value signifies the difference in OGG$_i$ across the two groups (Wilcoxon test). To generate a null distribution of p-values, we create 100 permutations of the distributions between groups A and B that randomly shuffles entries in the two groups. For all OGGs within the Spectral Tree, we compute a distribution of the most significant p-values across all permutations. The Q-value for a given OGG is computed as the number of p-values in the null distribution that are less than the reference p-value ($P_{ref}$)) relative to the total number of p-values in the null distribution $\left(N_{P_{perm}}\right)$ (equation).

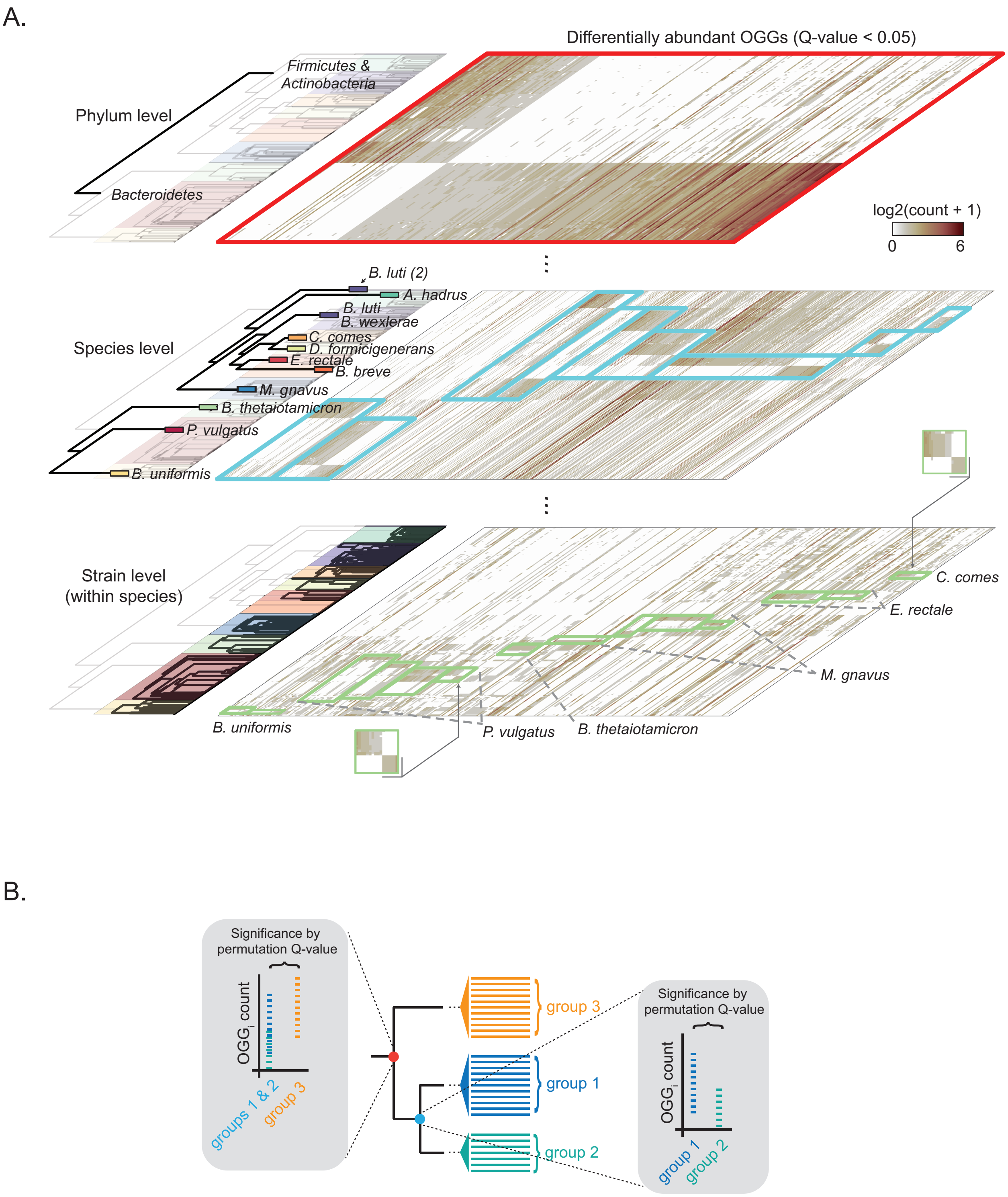

A.
Differentially abundant OGGs (Q-value < 0.05)
Phylum level
Firmicutes & Actinobacteria
Bacteroidetes
log2(count + 1)
0
6
Species level
B. luti (2)
A. hadrus
B. luti
B. wexlerae
C. comes
D. formicigenerans
E. rectale
B. breve
M. gnavus
B. thetaiotamicron
P. vulgatus
B. uniformis
Strain level (within species)
C. comes
E. rectale
M. gnavus
B. uniformis
P. vulgatus
B. thetaiotamicron
B.
Significance by permutation Q-value
OGG$_i$ count
groups 1 & 2
group 3
group 3
group 1
group 2
Significance by permutation Q-value
OGG$_i$ count
group 1
group 2

**Fig. S13.**
The Spectral Tree reveals a hierarchy of phylogeny through capturing nested genomic variation. **(A)** Set of layered dendrograms depict clusters of the Spectral Tree from shallow (top) to deeper (middle) to deepest (bottom). Hierarchy of clusters follow broad phylogenetic differences to subspecies phylogeny (see labels of each dendrogram). Heatmaps shows set of OGGs whose abundance is significantly different between clusters of strains defined at each layer (colored boxes). Pixel color indicates abundance of specific OGG. **(B)** The hierarchical nature of the Spectral Tree enables defining OGGs that are significantly different between daughter clusters that come from parent clusters. In this example, the red node splits group 3 from groups 1 and 2; the blue node splits group 1 from group 2. For each node, a significance test (Q-value) for every OGG is performed between the two groups (see **Fig. S12** for workflow of Q-value test).

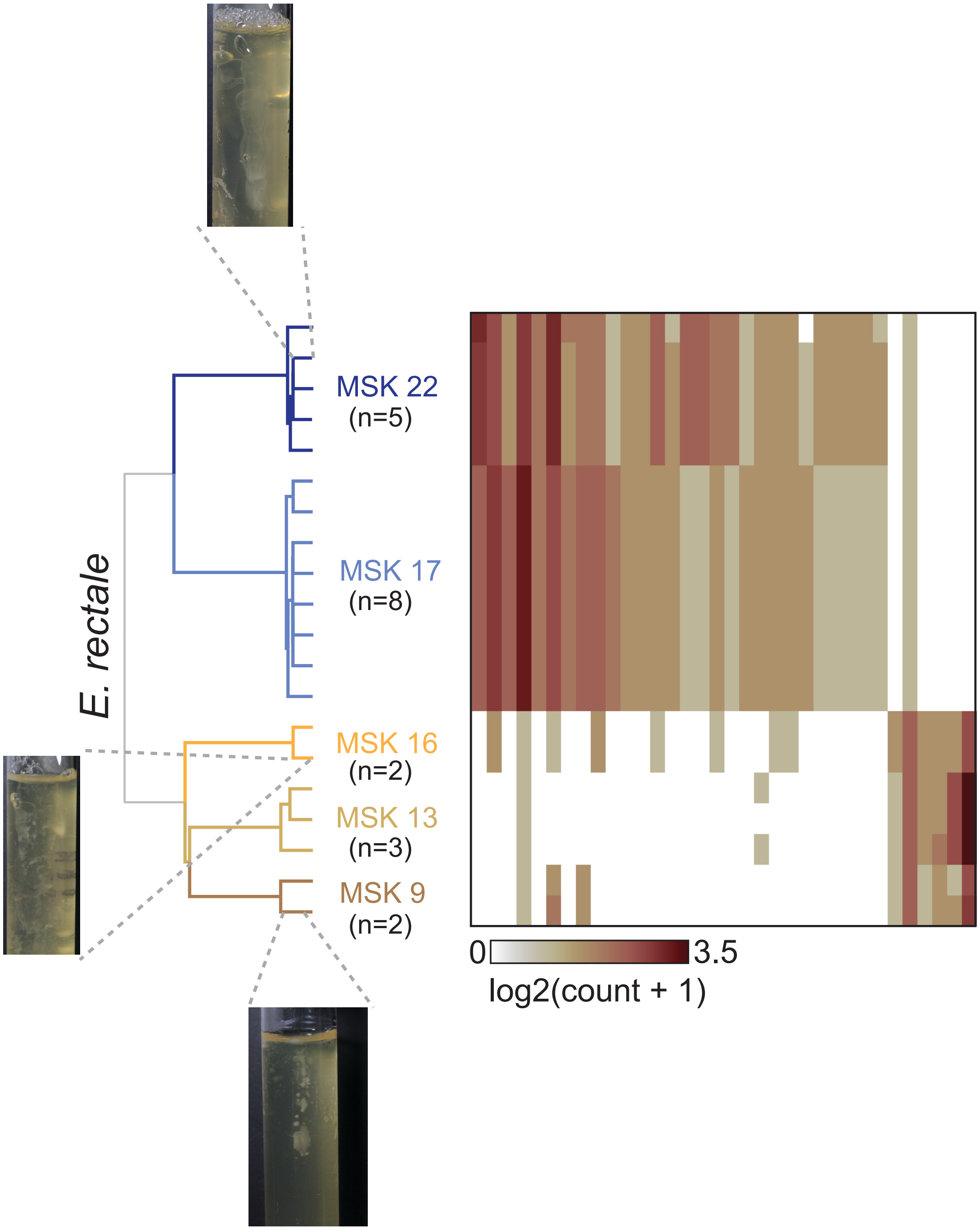


**Fig. S14.** Agar-based motility assay experiments for select *E. rectale* strains. Spectral tree and corresponding pattern of significantly differentially abundant OGG abundances from **Fig. 3A** shown.

A.

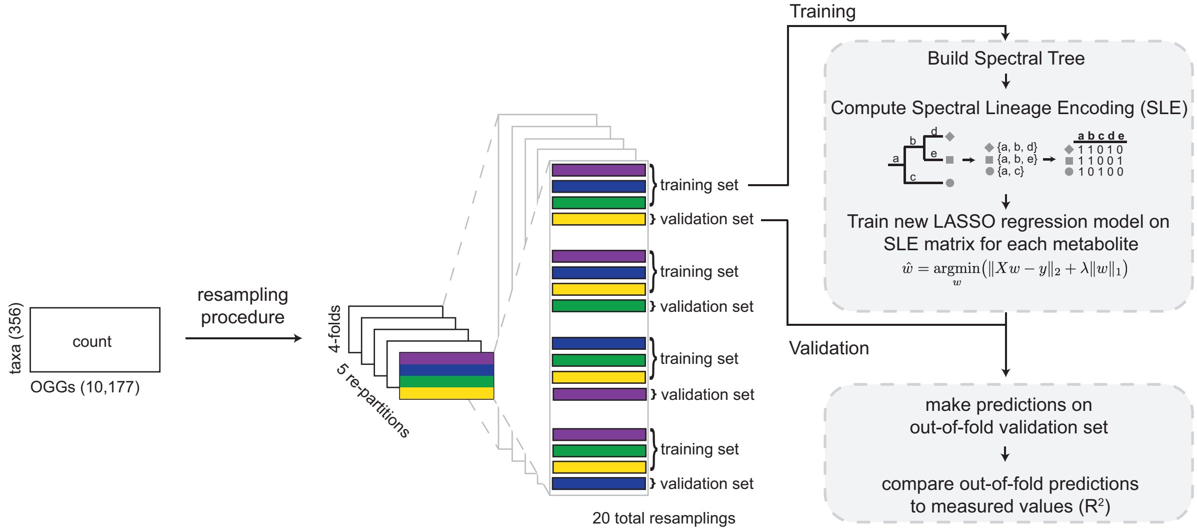


B.

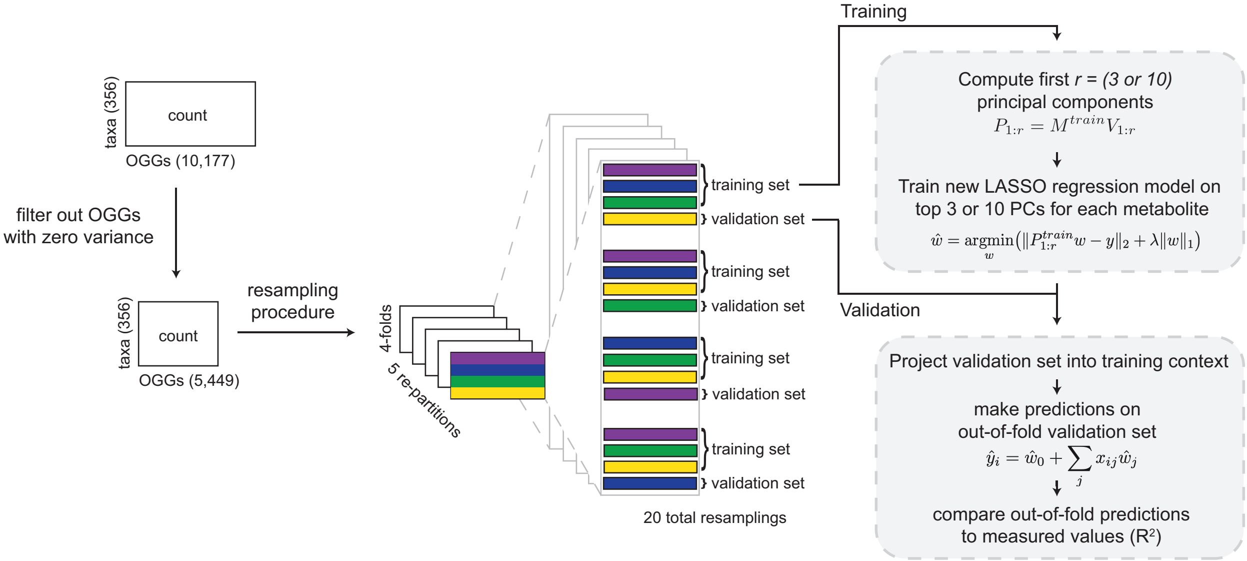


**Fig. S15.**
**(A)** Workflow for training SLE LASSO models. **(B)** Workflow for training LASSO models on top three or top ten principal components of training set.

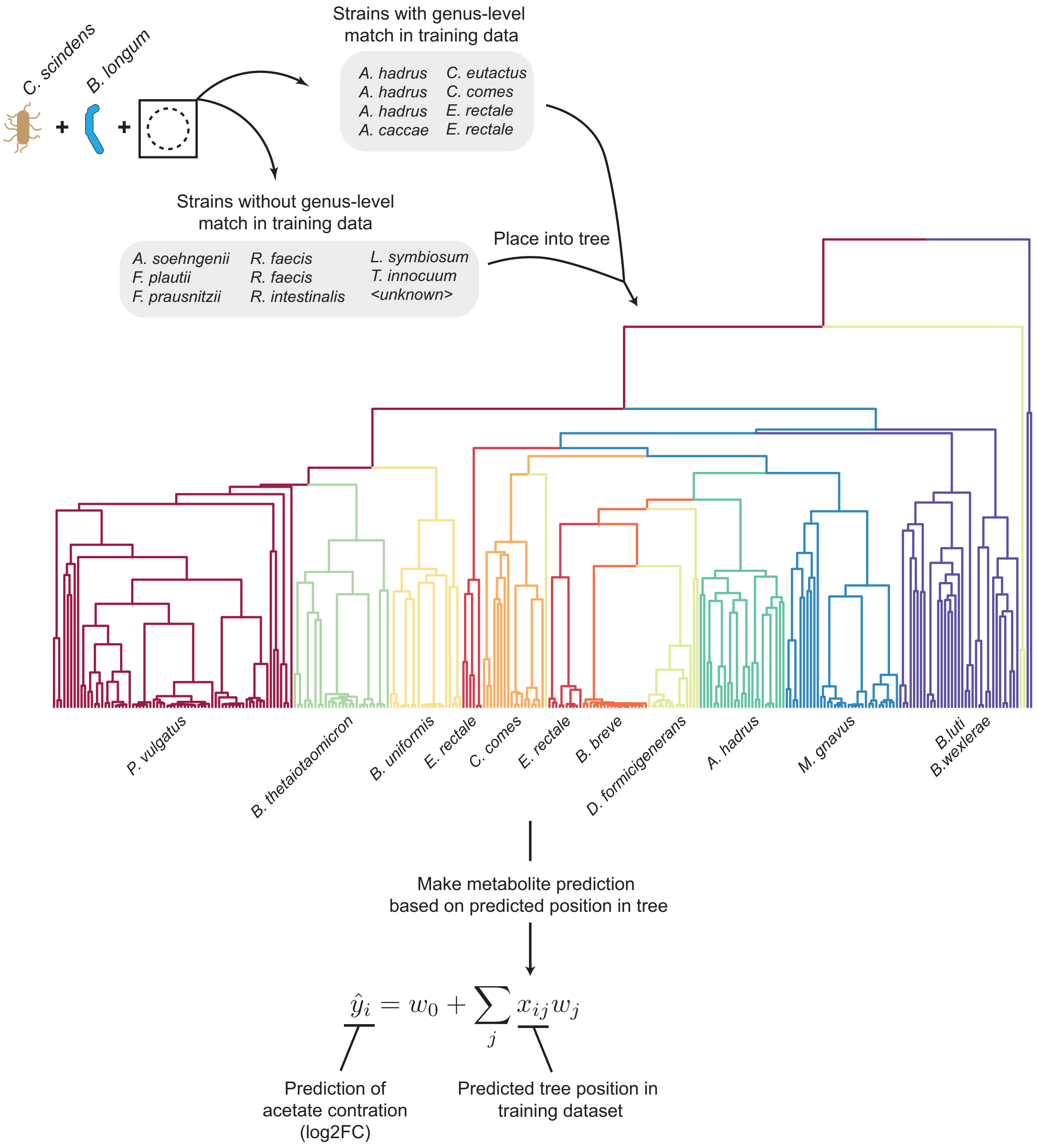


**Fig. S16.**
Phylogenetic distribution of seventeen strains used to create consortia shown in **Fig. 5** and workflow for predicting relative acetate concentration (log2FC) for each strain.

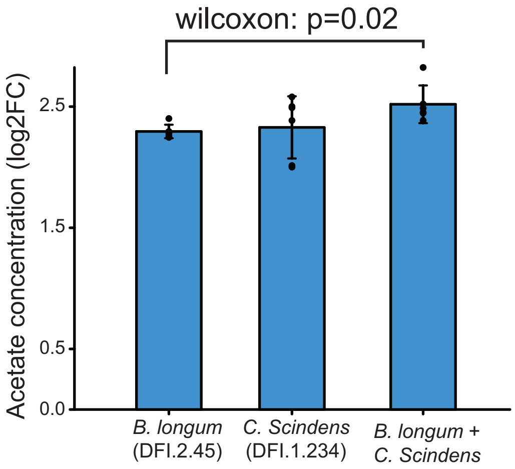


**Fig. S17.**
Bar graph of acetate concentration (log2FC as compared to empty BHIS media) for individual and combined strains used as the consortia background.

**Table S1. (separate file)**
Commensal strain bank described by strain ID and phylogenetic classification per NCBI and GTDB.

**Table S2. (separate file)**
Relative metabolite concentrations for Species with at least 10 representative strains in the commensal strain bank.

**Table S3. (separate file)**
Functional annotations of differentially abundant OGGs amongst strains in **Fig. 3D**.

**Table S4. (separate file)**
Predictions of relative metabolite concentrations from SLE-based LASSO models.

**Table S5. (separate file)**
Predicted acetate concentrations for each of the 17 strains in **Fig. 5**; acetate concentrations for each consortium described in **Fig. 5**.
